# Novel major loci shape habitat-associated flowering time variation in Yellowstone monkeyflowers

**DOI:** 10.1101/2025.06.30.662416

**Authors:** Colette S. Berg, Margaret Hendrick, Kory Kolis, Kipp Stebbins, Findley Finseth, Lila Fishman

## Abstract

- Plants harbor remarkable genetic diversity in flowering phenology, particularly in their responses to environmental cues such as photoperiod. Understanding the genetic basis of repeated evolution in flowering cues, which are key to reproduction, illuminates adaptation with gene flow and parallel evolution.
- We characterized variation in minimum critical daylength for flowering (MCD) in yellow monkeyflower (*Mimulus guttatus*) accessions from a geothermal soil mosaic in Yellowstone National Park, mapped loci underlying the most extreme MCD in focal thermal annuals, and investigated environmental variables shaping phenology in the field.
- Yellowstone monkeyflowers range in MCD from 12-15 hours, paralleling range-wide variation in *M. guttatus*; plants from thermal habitats flower under significantly shorter daylengths. Two QTLs govern the most extreme 12-hour MCD. Both contain candidates from gene families previously implicated in phenological evolution in monkeyflowers and other angiosperms, but the major loci appear novel. The frequency of 12-hour flowering across a microgeographic gradient is predicted by variation in soil temperature and the timing of dry-down.
- Adaptation to Yellowstone’s geothermal soil mosaic has generated dramatic evolution of flowering cues over short spatial scales. The genetic basis of 12-hour flowering does not indicate re-use of known *M. guttatus* alleles, but strong candidate genes nonetheless suggest molecular parallelism.

## Introduction

To successfully reproduce, angiosperms must flower when resources, mates, and pollinators are available in their local environment. Thus, flowering phenology often varies widely across a species range, due both to direct environmental effects and population differentiation in genetically-encoded cues. The latter include external factors such as daylength (e.g., Blackman *et al*., 2011) and vernalization (Ratcliffe *et al*., 2003; Yan *et al*., 2004), as well as endogenous factors such as age or size (Amasino, 2010). Critical photoperiod cues, such as the minimum daylength required to initiate floral buds for long-day plants allow plants to anticipate future growing conditions in advance. Thus, populations from lower latitudes (Friedman & Willis, 2013; Ågren *et al*., 2017) or altitudes (Lewandowska-Sabat *et al*., 2017) tend to have shorter daylength requirements that allow spring flowering prior to summer drought. Such local fine-tuning of cues facilitates temporal escape from detrimental environmental conditions and is a key component of local adaptation and potentially reproductive isolation in species with broad geographic ranges. However, phenology may also vary adaptively over much shorter spatial scales, as microgeographic soil mosaics that affect growing season length can create divergent selection within a single climate (Savolainen *et al*., 2006; Antonovics, 2006; Selby & Willis, 2018; Sianta & Kay, 2021). Understanding the ecological, genetic, and evolutionary basis of such phenological variation, and how it influences patterns of gene flow, illuminates a major mechanism of plant adaptation and its consequences for speciation.

Abundant flowering time variation across populations provides raises fundamental questions about the genetic architecture of adaptive divergence (Gaudinier & Blackman, 2020). Repeated shifts in photoperiod and temperature cues provide the opportunity to ask whether parallel phenotypic evolution occurs through novel genetic changes (e.g., Fernández-Ortuño *et al*., 2008), re-sampling of shared standing variation (e.g., Jones et al., 2012; Nelson & Cresko, 2018), or convergent *de novo* mutations to the same set of loci (e.g., Chan *et al*., 2010; Rosenblum *et al*., 2010). Compared to quantitative traits such as flower size, flowering time variation appears to be constrained at the molecular level. In particular, variants at genes related to the florigen Flowering Locus T (FT) influence phenological divergence in diverse angiosperms (reviewed in Pin & Nilsson, 2012; Putterill & Varkonyi-Gasic, 2016; Gaudinier & Blackman, 2020). Further, with the exception of maize (Buckler *et al*., 2009), even dramatic differences in minimum daylength requirements often have simple genetic architecture (Salomé *et al*., 2011; Navarro *et al*., 2011; Koskela *et al*., 2012; Fishman *et al*., 2014; Weller & Ortega, 2015). This suggests a finite number of targets for selection within the flowering pathways, but the same basic players may be re-configured in different taxa (Salomé *et al*., 2011; Fournier-Level *et al*., 2022). Furthermore, because similar selection on flowering time can occur over both microgeographic (10-100s of meters) and latitudinal (100s of kilometers) spatial scales, understanding the genetics of daylength requirements in both contexts provides insight into the predictability and repeatability of evolution.

Here, we investigate the patterns, mechanisms and drivers of genetic variation in daylength requirements for flowering in *Mimulus guttatus* (yellow monkeyflower) from a dramatic microgeographic soil mosaic created by geothermal inputs in Yellowstone National Park. Yellow monkeyflowers of the *M. guttatus* species complex are common in seasonally wet soils across western North America (Wu et al. 2008), and exhibit tremendous variation in life history (e.g., annual vs. perennial; Lowry & Willis, 2010; Friedman *et al*., 2015), mating system (Fishman *et al*., 2002), and daylength requirements for flowering (8-16 hours) (Friedman & Willis, 2013) across a geographic range extending from Alaska to Mexico and from the Pacific Coast to the Rockies. Phenotypic variation in these traits tends to be correlated at both local (Kolis *et al*., 2022) and latitudinal scales (Hall & Willis, 2006; Lowry & Willis, 2010; Friedman *et al*., 2015). This is because yellow monkeyflowers are highly adaptable (Wu *et al*., 2008) but cannot tolerate drought. Thus, summer-dry habitats (e.g., snowfields, seeps) foster annual ecotypes, while those with soil moisture throughout the summer (e.g., bogs, coastal springs, permanent riverbanks) harbor perennials. Within each group, daylength requirements further vary with altitude and latitude (Fishman *et al*., 2014; Friedman *et al*., 2015; Kooyers *et al*., 2015) and self-pollination and early spring flowering has evolved in annuals adapted to the most ephemerally wet sites (Fishman *et al*., 2014; Kenney & Sweigart, 2016; Mantel & Sweigart, 2019). Remarkably, the interspecific transition between <12-hour (selfer *M. nasutus*; early March) and >16hr (Iron Mountain high elevation annual *M. guttatus*; late June) daylength requirements for flowering are entirely explained by two major loci on Chromosome (Chr) 7 and Chr 8 in both allopatric and sympatric contexts (Fishman *et al*., 2014). A similar shift between inland annuals and coastal perennials is largely explained by a shared QTL on the end of Chr 8 (Hall *et al*., 2010; Friedman & Willis, 2013), plus the major life-history diagnostic inversion (*inv8*; (Hall *et al*., 2006; Lowry & Willis, 2010) on the other end of that chromosome. The partial overlap in QTLs suggests that variation in daylength requirements for flowering may have a predictable genetic architecture across the *M. guttatus* complex. Furthermore, the presence of functional candidates homologous to *Arabidopsis FLOWERING LOCUS T* (*FT*) and *SHORT VEGETATIVE PHASE* flowering time genes, respectively, within the Chr 7 and Chr 8 QTL regions (Fishman *et al*., 2014), suggests conservation of key components of the critical photoperiod pathway across diverse plants.

Here, we characterize the diversity of daylength requirements for flowering in yellow monkeyflower (*M. guttatus)* populations growing across Yellowstone National Park’s (YNP) geothermal soil mosaic, as well as the genetic basis and environmental correlates of one extreme microgeographic (<100m) transition paralleling the *M. guttatus* - *M. nasutus* interspecific divergence. Yellow monkeyflowers are one of only a handful of angiosperm species able to live in the extreme soils around Yellowstone geothermal features, which generate a meter-scale habitat mosaic that recapitulate diverse conditions typically found varying at continental scale. YNP geothermal soils are generally lethally hot and dry by early summer but winter/spring snowmelt on hot soils creates a near-ground micro-climate suitable for growth and flowering. Thus, thermal *M. guttatus* populations have locally evolved annuality, increased self-fertilization ability, and spring flowering to take advantage of narrow, temporally-defined habitat (Rice, 1973; Lekberg *et al*., 2012; Kolis *et al*., 2022). At its most extreme, thermal adaptation necessitates a shift to <12-hour daylength to initiate flowering under the short days of spring (i.e., March) when snow is meters-deep on nonthermal soils but thermal crusts are snow-free and still habitable (Lekberg *et al*., 2012). However, it is unclear whether and how daylength requirements vary across the diverse mosaic of thermally-influenced habitats, some of which remain wet and warm into midsummer (Kolis *et al*., 2022). Intriguingly, thermal annual YNP *M. guttatus* (as well as nonthermal perennials) have the perennial orientation of the widespread *inv8* inversion governing life history throughout most of the range, suggesting a novel genetic basis for the life history shift (and possibly phenological evolution) in YNP (Kolis *et al*., 2022). Furthermore, high connectivity between thermal annula and nonthermal perennial habitats in YNP (Kolis *et al*., 2022) provide a unique setting to assess the genetic basis of microgeographic variation in flowering cues in the context of broader variation across the *M. guttatus* species complex.

First, to characterize how YNP’s geothermal mosaic has shaped flowering cues and test the hypothesis that thermal annual populations will generally flower under shorter days than non-thermal neighbors, we assess flowering in a set of YNP-wide inbred lines at four daylengths (12,13 14, 15 hours). We map the genetic basis of extreme flowering time divergence in F_2_ hybrids between one thermal annual (12-hour daylength requirement) and a nearby (< 200m) nonthermal perennial (15-hour daylength requirement). We predict that this shift will be under the control of a small number of genomic regions, as with parallel transitions. We further refine the pool of candidate genes and variants underlying our QTLs by simulating a threshold for the plausible frequency of the causal variant in whole genome sequenced inbred lines. This finer-mapping allows us to ask about parallelism at the scale of many-gene QTLs and candidates within them. Finally, we assess variation in daylength requirements (12-hour flowering vs. not), natural phenology, and environmental variables along a transect between the extreme thermal annual and nonthermal perennial populations represented by our QTL mapping parents. This allows us to ask how well genetic variation in daylength requirements mirrors a gradient in wild phenology and whether soil/air temperature and/or the timing of soil dry-down predicts either. Together, these experiments illustrate how soil mosaics drive genetic divergence in a key plant reproductive trait, dissect the genetic and ecological causes of an extreme shift in minimum critical daylength requirements in a microgeographic context, and contribute to understanding of how gene flow and natural selection shape the genetics of local adaptation.

## Methods

### Study System

*Mimulus guttatus* Fisch. Ex. DC (Phrymaceae) (also known as *Erythranthe guttata*, see Lowry *et al*., 2019; Nesom *et al*., 2019) is an herbaceous plant common throughout western North America in diverse wet soil habitats. In Yellowstone National Park (YNP), *M. guttatus* grows in a complex mosaic of geothermally-influenced soils, ranging from highly thermal crusts to nonthermal bogs. *M. guttatus* bloom in spring on thermal soils, when geothermal inputs melt snow and create a temperate ground-level climate, and are annual, dying in summer when thermal soils are lethally hot and dry (Lekberg *et al*., 2012; Kolis *et al*., 2022). In contrast, *M. guttatus* in nonthermal habitats are genetically perennial, producing stolons for overwintering beneath the snow and flowering in mid-late summer (Lekberg *et al*., 2012; Kolis *et al*., 2022). One extreme thermal annual population from Agrostis Headquarters, AHQT (Fig. 1), is genetically distinct from all other YNP *M. guttatus* populations (F_ST_ = 0.279 - 0.348, Kolis *et al*., 2022). AHQT plants have a 12-hour daylength requirement, while plants from the nearby nonthermal bog population AHQN (only ∼200 meters away) only flowers at daylengths >15 hours (Lekberg *et al*., 2012; Hendrick *et al*., 2016; Kolis *et al*., 2022). AHQT appears exceptional in its bottlenecking and isolation; all other adjacent thermal annual and nonthermal perennial populations throughout YNP are relatively undifferentiated genetically (F_ST_ range = 0.04-0.07, Kolis *et al*., 2022), though they are often almost as different morphologically and phenologically as the AHQ extremes.

**Figure 1.**
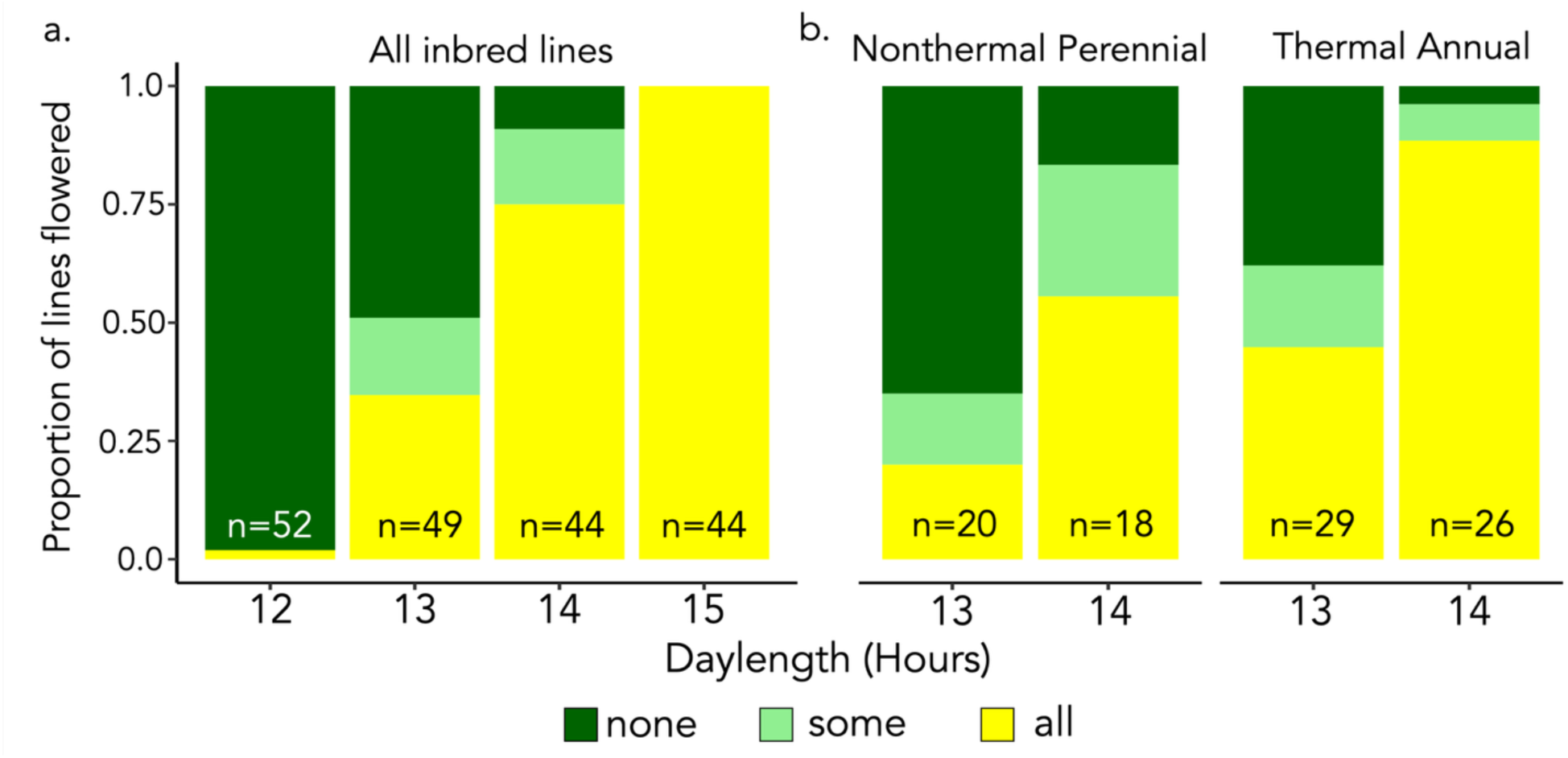
Critical photoperiod varies in Yellowstone National Park *Mimulus guttatus* inbred lines from thermal and nonthermal habitats. a. Proportion of flowering in inbred lines under 12, 13, 14, and 15 hours of daylength (1 – 4 reps per line). b. Proportion of flowering for in inbred lines under 13 and 14 hours of daylength, split by habitat type (nonthermal perennial or thermal annual). Dark green = never flowered, light green = some flowered, yellow = always flowered.

### Parkwide survey of daylength requirements for flowering

To characterize genetic variation in the daylength required for flowering across the YNP habitat matrix, we generated inbred lines (n = 57, Table S1) from a subset of accessions previously characterized (Kolis *et al*., 2022). Lines were made by four generations of self-fertilization with single seed descent. Throughout line generation and in all other experimental grow-outs described below, plants were grown in Sunshine Soil Mix #4 (SunGro Horticulture, Agawam, MA, USA) with Supplementary lighting, daily bottom-watering to saturation, and monthly fertilization with 20:20:20 N:P:K fertilizer (Peters Professional, Tel Aviv, Israel). We tested each line for flowering under 12-, 13-, 14- and 15-hour daylengths (20℃ days, 15℃ nights) in identical PCG40 growth chambers (Percival Scientific, Perry, IA, USA) with line positions scrambled in 96-well cell-trays (n = 1 - 4 reps per line per chamber; total N = 446). Each plant was scored as bolted (1) vs. not (0) after 12 weeks. Lines were categorized as “fully flowering”, “not flowering”, or “mixed” at each daylength, and we compared the proportion of fully flowering lines from each habitat for each daylength treatment with chi-square tests.

### QTL Mapping the genetic basis of flowering under 12-hour days

We characterized the genetic basis of the most extreme divergence in daylength requirements using an F_2_ hybrid mapping population between inbred lines from adjacent (< 200 m) thermal annual (AHQT; 12-hour daylength requirement) and nonthermal perennial (AHQN; 15-hour daylength requirement) populations. Lines were generated through > 4 generations of self-fertilization with single-seed descent and crossed to generate an F_1_, which was then self-fertilized to generate F_2_s. F_2_ hybrids (n = 288), plus AHQT (n = 1), AHQN (n = 1), and F_1_ (n = 2) controls, were grown under 12-hour days. Individuals were scored as bolted (1) vs. not (0) after 12 weeks. To define parental genotypes, we used whole-genome sequence from the parents: for the AHQT parent we used existing whole genome sequence (Brandvain *et al*., 2014); whole genome sequence from the AHQN parent was prepped using an NEBNext illumina prep kit and sequenced by HudsonAlpha on a NovaSeq 6000. We aligned AHQT and AHQN to the IM62v2 reference genome using bwa mem (Li, 2013) and called SNPs using GATK-4.0 (McKenna *et al*., 2010) with default parameters.

For ddRAD genotyping, we extracted F_2_ and control DNA with a CTAB-chloroform protocol in 96-well plates (https://dx.doi.org/10.17504/protocols.io.bgv6jw9e), generated ddRADseq libraries using a modified BestRAD protocol (Nelson, 2019), and sequenced the F_2_s on 37.5% of a HiSeq4000 lane at GC3F (University of Oregon). Reads were demultiplexed using custom python scripts (dx.doi.org/10.17504/protocols.io.bjnbkman) and adapter sequences were removed with Trimmomatic (Bolger *et al*., 2014). Reads were aligned to the *M. guttatus* IM62v2 reference genome (Goodstein *et al*., 2012) using bwa mem with default parameters (Li, 2013) and GATK-4.0 (McKenna *et al*., 2010) for indel realignment. SNPs were called with GATK-4.0 (expected heterozygosity = .05, all other parameters default, McKenna *et al*., 2010). We retained F_2_’s with > 150,000 reads (n = 196), which had an average coverage > 4, then filtered to fully informative sites with fixed differences between the parental AHQT and AHQN lines. SNPs were filtered to biallelic sites using vcftools (Danecek *et al*., 2011) to a minimum depth of 3, maximum missing of 0.5, minimum quality of 29, and minor allele frequency of 0.05. We retained one SNP per read, then manually filtered to sites that fit transmission ratio expectations for an F_2_ mapping population (homozygous reference frequency > 0.05 and < 0.4, homozygous alternate genotype > 0.05 and < 0.35). Because AHQT and AHQN lines are derived from populations < 200m apart, there were swathes of the genome that were not differentiated; we did not recover informative markers on 11-13 Mb of scaffold 1, 0-8 Mb of scaffold 9, 5.3 – 17.1 Mb of scaffold 11, or 0-4 Mb of scaffold 14. The filtered genotype dataset consisted of 1422 markers.

We generated a linkage map in lepMap3 (Rastas, 2017), using AHQT and AHQN parental genotypes for phasing (command = ParentCall2). Linkage groups were separated (command = SeparateChromosomes2, LOD limit = 12, theta = 0.05), and markers were ordered with a Kosambi mapping function (command = OrderMarkers2). LepMap3 produced a map with 1273 markers spanning 929 cM and corresponding to 247Mb of the *M. guttatus* IM62v2 reference genome (280 Mb total). We scanned for QTLs underlying 12-hour flowering with the composite interval mapping (CIM) function (window size = 10 cM, 5 background markers) in QTLCartographer (Basten *et al., 2006*). A genome-wide significance threshold (LOD = 3.7) was calculated from 1000 permutations of the phenotypes vs. genotypes. We tested for interaction between the most significant mapped QTLs using logistic regression in JMP (Jones & Sall, 2011)

### Identifying candidate variants underlying AHQT’s extreme 12-hour daylength requirement

To identify candidate variants unique to AHQT (the only inbred line which flowered under 12-hour days), we quantified the frequency of reference (AHQT) alleles within the QTL regions in 44 inbred lines that did not flower under those conditions. Inbred line libraries were prepped with Illumina tagmentation kits (Illumina, Inc., San Diego, CA) and sequenced as part of NovaSeq 6000 lane (150bp, paired end reads). We trimmed reads with Trimmomatic (Bolger *et al*., 2014) and aligned all inbred lines (as well as existing AHQT sequence) to the AHQTv1 reference genome (RIfkin *et al. in prep*), using bwa mem with default parameters (Li, 2013). We called SNPs using GATK-4.0 (expected heterozygosity = 0.05, all other parameters default), and filtered with vcftools (Danecek *et al*., 2011) to sites with a minimum depth of 3, maximum missing of 0.2, and minimum quality of 29.

To narrow down to potentially causal SNPs within the strongest 12-hour flowering QTL, we used a simple simulation, integrating the findings from the inbred line daylength screening and the QTL map. Because only one line, AHQT, flowered under 12-hour days in the inbred line screening, and AHQT is the reference genome line, the causal allele for 12-hour flowering should be present in the AHQT line but rare or absent in the other inbred lines. Furthermore, AHQN (the non-flowering parent in the F_2_ mapping population) should carry the alternate allele at the causal site. To set a specific threshold for the predicted frequency of the causal allele in the non-flowering inbred lines, we calculated the 95% confidence intervals for the likelihood of flowering at each genotype in the QTL map, conservatively assuming that the weaker QTL is fixed for the non-flowering allele. The inbred lines are four generations inbred, so there is a nearly 1:1 relationship between the frequency of the reference (T) allele and the frequency of the TT genotype (t = 0.999, p = <0.0001). Therefore, we predicted the expected frequency of flowering in inbred line populations at reference allele frequencies ranging from 0 – 1 at intervals of 0.05, multiplying each homozygous genotype frequency (TT or NN) by the frequency at which an individual with that genotype flowered under 12-hour days in the QTL map, and adding both terms to get the total flowering frequency predicted in the inbred lines. With these predictions, we identified a plausible threshold for the frequency of the causal 12-hour flowering allele in the non-flowering inbred lines. Because our 95% confidence intervals for flowering frequency at each genotype are wide and we assumed that the weaker QTL is fixed for the alternate (non-flowering) allele, our threshold is relatively generous: it may include variants that are highly differentiated but not causal, but it is unlikely to erroneously exclude the causal variant. We defined candidate SNPs as those in which AHQT has the reference allele, AHQN has the alternate allele, and the alternate allele frequency is > 0.8 in the non-flowering inbred lines.

To identify *a priori* candidate genes within *sd6*, we extracted the *Arabidopsis thaliana* homolog gene IDs from the MgAHQTv1 reference genome annotation (Goodstein *et al*., 2012) and searched for the functional categories associated with these genes using the TAIR Functional Categorization tool (Rhee *et al*., 2003). We identified genes whose *A. thaliana* homologs have GO terms corresponding to flowering, photoperiod, plant organ morphogenesis, and meristem development as *a priori* candidates for shaping flowering time. Because environmental cueing to flower involves integration of signals from multiple plant pathways mediated by plant hormones (Izawa, 2021), we identified hormone-related genes with *sd6*. Within the AHQTv1 annotation, we searched for gene descriptions that included the major plant hormones auxin, gibberellin, cytokinin, ethylene, abscisic acid, brassinosteroid, salicylic acid, jasmonate, and strigolactone (Waadt, 2020). For the 31 remaining genes that had GO terms but no *Arabidopsis* homologs, we searched the AHQTv1 annotation for GO terms related to flower development, photoperiodism, flower morphogenesis, flower formation, meristem development, maintenance of meristem identity, regulation of floral meristem growth, meristem determinacy, and vegetative to reproductive meristem transition. For the 7 genes which had no GO terms or *Arabidopsis* homologs, we used the Phytozome gene def-lines if available (Goodstein *et al*., 2012).

The weaker effects and broad span of the second mapped QTL, *sd11*, prevented a similar modeling approach. However, this region contains a cluster of *a priori* candidate genes for phenological traits, in the *MADS AFFECTING FLOWERING* (MAF) family. To quantify copy number variants in this cluster, we median-standardized the coverage for AHQT and AHQN, multiplying the coverage at each read by the genome-wide genic coverage median for that individual and subtracting the genome-wide median absolute deviation. With this standardization method, negative coverage values can occur where coverage is lower than the median for that individual (AHQT or AHQN), but still present. Deleted genes have negative standardized values, and very little variance.

### Investigating fine-scale variation in daylength requirements, flowering phenology, and environmental factors at AHQ

To understand fine scale variation in daylength requirements for flowering, we focused on a microgeographic transect spanning the AHQT thermal annual and AHQN nonthermal perennial extremes as well as intervening thermal sites (AHQCanyon) (Hendrick *et al*., 2016), We characterized the ability to flower under 12-hour days in plants grown from wild-collected seeds from AHQT (n = 107 individuals in 20 maternal families), AHQN (N = 18 individuals in 7 families) and AHQCanyon (N = 698 individuals in 78 families) under 12-hour days.

To compare patterns of natural flowering phenology and site dry-down (i.e. *Mimulus* death) to daylength requirements for flowering along the AHQ transect, we leveraged phenological and environmental data from a series of 0.5m x 0.5m quadrats monitored in the 2010-2011 growing seasons (Hendrick *et al*., 2016). Each quadrat (n = 10 total) was photographed at intervals (as vehicle access allowed) and phenology then scored at 13 time points from April 2010 – April 2011 (Table S3). For each photograph at each time point, we counted the number of open flowers and scored the quadrant as “pre-flowering” (0 open flowers), “flowering” (≥ 1 open flower), or “post-flowering” (all individuals senesced). Quadrats in peak flower (maximum number of flowers open) by late April were scored as “early”; quadrats with peak flowering in mid-June were scored as “late”. Each quadrat was centered around a pair of aboveground (leaf height) and belowground (root height) temperature dataloggers (DS1921G-F5 Thermochron iButton; Embedded Data Systems, Lawrenceburg, KY, USA) recording plant and soil temperatures every 4 hours, as previously described in (Lekberg *et al*., 2012). In addition, to capture the soil drying that leads to *M. guttatus* death in thermal annual soils (Kolis *et al*., 2022), we characterized soil water using the wooden toothpick method (Martin, 2004). For each sampling point, a toothpick was buried for 24 hours near each datalogger, collected into a 2ml tube (screw-cap lid with O-ring), tightly sealed in the field, and then stored in a −80 freezer. The tubes (with toothpicks in place) were then weighed on a microbalance to nearest 0.01mg, opened and dried in a drying oven for 48 hours, then reweighed on the same balance. Relative soil moisture was calculated as the difference between the pre- and post-drying weights for each individual toothpick; these values were standardized with water-mass estimates from unburied toothpicks (0 = maximally dry; n = 10) and toothpicks soaked in water for 24 hours (1= maximally wet; n = 9).

To assess how well environmental variables predict flowering phenology in AHQT and AHQCanyon, we conducted an ordinal logistic regression in JMPv18.0.1 (SAS Institute, Cary, NC) with flowering phase (pre-flowering, flowering, or post-flowering) as the response variable and soil moisture, maximum soil temperature, and maximum plant temperature as the explanatory variables. Additionally, we compared 4-PM plant and soil temperatures between early and late flowering quadrats at AHQ at 4PM (the warmest part of the day) on the peak flowering date in lower AHQCanyon.

## Results

### *M. guttatus* daylength requirements vary widely but predictably across the Yellowstone geothermal mosaic

Minimum daylength required for flowering for flowering ranged from 12 to15 hours in YNP *M. guttatus* accessions, recapitulating nearly range-wide variation within the microgeographic geothermal soil mosaic (Fig. 1). Only the AHQT line from the focal extreme thermal annual population flowered under 12-hour days, and all lines flowered under 15-hour days. Thermal annual lines were more likely to flower than nonthermal perennials at intermediate daylengths: 48% of thermal annuals and 20% of nonthermal perennial lines flowered fully under 13-hour days, although the relationship was not quite significant (p = 0.067), and 92% of thermal annuals and 55% of nonthermal perennials flowered under 14-hour days (p = 0.005) (Fig. 1).

Two loci explain microgeographic divergence in minimum daylength requirements for flowering (12-hour vs. 15-hour)

Two QTLs, *short day 6* (*sd6*, LOD = 28.8) and *short day 11* (*sd11*, LOD = 6.7), govern 12-hour flowering in the AHQT x AHQN F_2_ mapping population (Fig. 2a). The flowering-permissive AHQT allele at *sd6* shows partial dominance (94% of homozygotes and 72% of heterozygotes flower under 12-hour days, Fig. 2b). *sd6* has a stronger effect on flowering than *sd11* (*sd6* R^2^ = 0.41, *sd11* R^2^ = 0.08), and the two loci do not exhibit epistasis (genotypic interaction *p* = 0.27 in logistic regression). The confidence interval (1.5 LOD drop) for *sd6* spans a 106-gene region near the end of Chr 6 (6.25cM, 260 kb) and *sd11* spans 663 genes (14.14cM, 18 Mb) on Chr 11 (Fig 2a). The stronger QTL, *sd6* is in a region of high marker and gene density, while *sd11* spans the structurally complex MDL11 locus, which was mis-ordered in the IM62v2 reference genome (Finseth *et al*., 2021).

**Figure 2.**
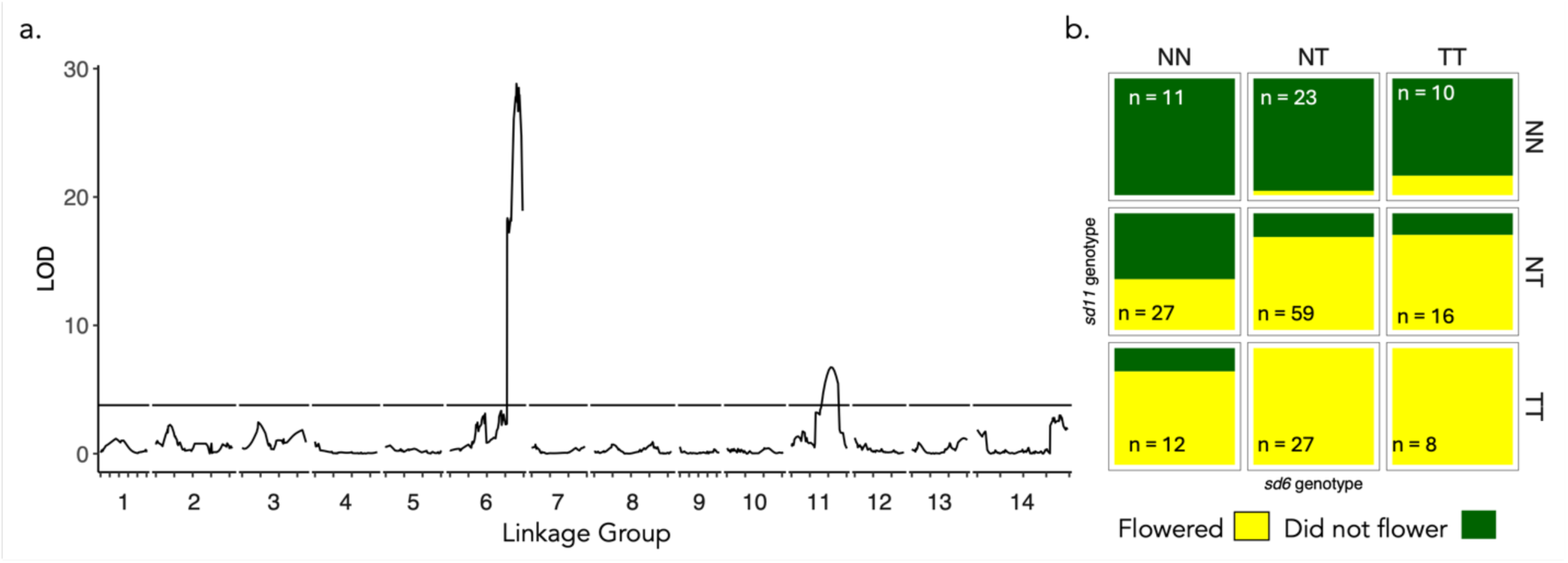
Major QTLs govern 12-hour flowering in Yellowstone *Mimulus guttatus*. a. Quantitative trait locus (QTL) mapping of 12-hour critical photoperiod for flowering in Yellowstone Mimulus guttatus (AHQT x AHQN F2 hybrids; permutation-based LOD threshold = 3.7). b. Proportion of F2 hybrids that flowered in the 9 genotype combinations for the two significant QTLs, *sd6* and *sd11* (sample sizes are listed in each block). N = non-flowering AHQN alleles, T = flowering AHQT alleles. Yellow = flowered under 12-hour days, green = not did not flower.

Both QTL regions contain strong *a priori* candidate genes for divergence in daylength requirements for flowering (Fig. 3, Fig. S2). The *sd6* LOD peak is within 20kb of a *FLOWERING LOCUS T* (FT)-like gene (MgAHQT1.06G203400); FT is the florigen-encoding gene considered a universal promoter of flowering in angiosperms (Wigge, 2011; Pin & Nilsson, 2012). The FT-like gene is a strong *a priori* candidate due to its role in flowering time in many systems including *M. guttatus* (Fishman *et al*., 2014). Beyond the *FT*, there were 3 additional *a priori* candidate genes within *sd6* whose *Arabidopsis* homologs are associated with flowering, meristem identity, or floral organ morphogenesis. *APETALA-LIKE* 2 (AHQTv1.06G203900) is similar to an *Arabidopsis* floral homeotic gene that controls the establishment of flower meristem identity among other important plant processes (Okamuro *et al*., 1997; Huang *et al*., 2017); *AP2-*like genes influence flower development in other systems as well, including maize (Zhuang *et al*., 2010) and petunias (Maes *et al*., 2001). The second-to-last gene before the chromosome 6 telomere is an auxin efflux carrier family gene (Fig 4); auxin is an important plant hormone which mediates diverse plant processes and is especially critical for growth (Woodward, 2005). Finally, the last gene before the Chr 6 telomere (Fig 4) is a homolog of the *Arabidopsis* gene *CLAVATA2*, a regulator of meristem and organ development (Kayes & Clark, 1998). The full list of *sd6* genes and their annotated functions is in Table S2. The broad *sd11* QTL most notably contains a cluster of 12 genes in the *MADS AFFECTING FLOWERING (MAF)* family of transcription factors, which interact with *FT*s to promote or repress flowering in a variety of systems (Yan *et al*., 2004; Deng *et al*., 2011).

**Figure 3.**
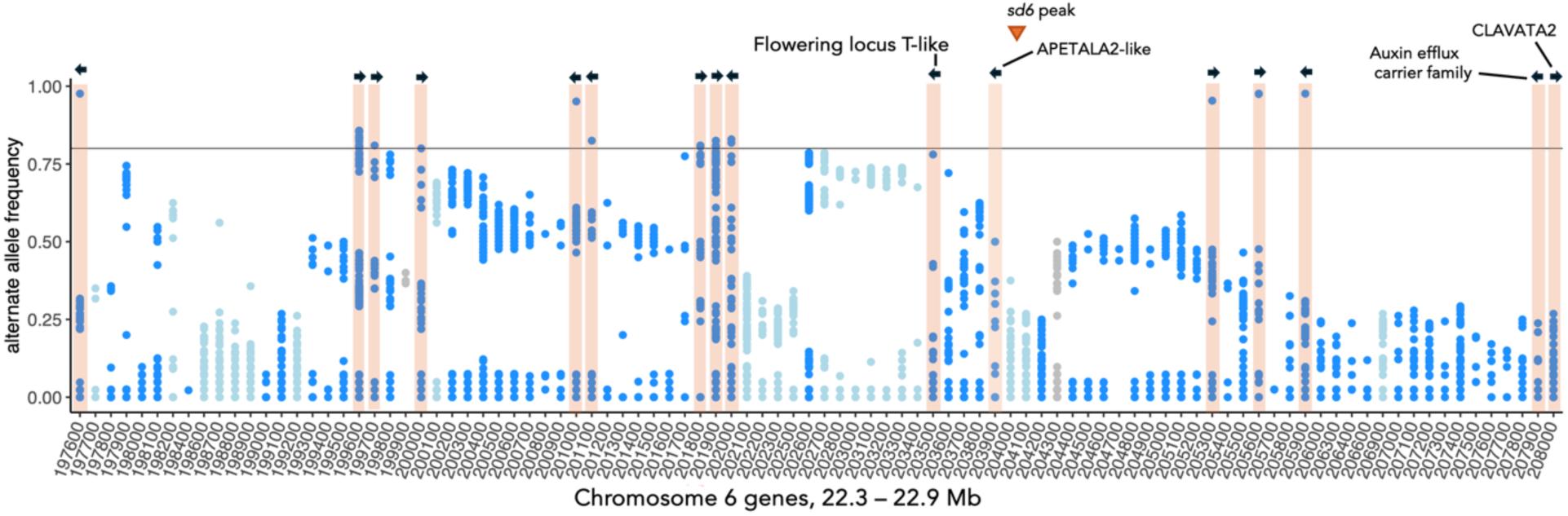
Candidate variants within *sd6*, the major QTL governing 12-hour flowering. Alternate (non-AHQT) allele frequency in non-12-hour flowering inbred lines, within each gene in the *sd6* QTL. Points with alternate allele frequency above 0.8 are candidate variants; genes with candidate variants are highlighted and the arrows represent gene direction (all *sd6* genes and annotations in Table S2). *A priori* candidates for involvement in flowering time are highlighted and labeled. The *sd6* peak is marked with an arrow.

**Figure 4.**
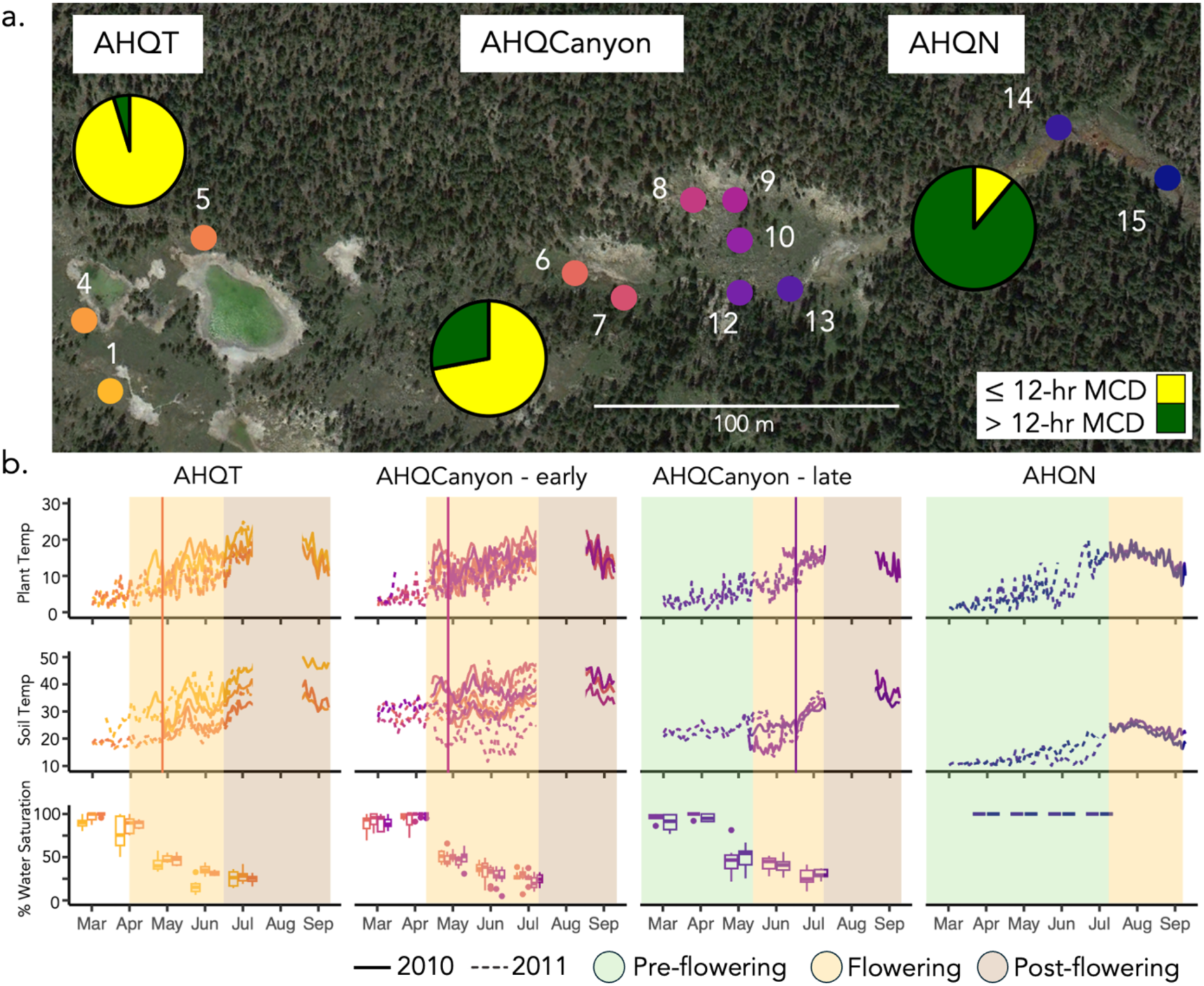
Fine-scale flowering phenology and geothermal variation in the extreme AHQ habitat. a. Aerial view of AHQT, AHQCanyon, and AHQN habitats. Pie charts represent the proportion of wild-collected seeds from each habitat that flowered in growth chambers under 12-hour days (yellow = flowered, green = did not flower.) b. Smoothed plant (top) and soil (middle) temperature data from dataloggers in quadrats at AHQT, early-flowering AHQCanyon, late-flowering AHQCanyon, and AHQN quadrats from March to August (solid line = 2010, dashed line = 2011). The pre-flowering window, if captured, is highlighted green; the flowering window is highlighted yellow, and the post-flowering (senescence) window is highlighted brown. Peak flowering at AHQT and AHQCanyon is indicated with a vertical line; our phenology photographs did not capture peak flowering at AHQN. Bottom: percent water saturation of soil at all quadrats at multiple time points, combined for 2010 and 2011.

### Population genomics refines candidate causal variants within *sd6* and *sd11* QTLs

None of the inbred lines from throughout YNP (n = 45) except AHQT flowered under 12-hour days (95% CI = 0 – 0.1). We used the frequency at which *sd6* TT and NN F_2_’s flowered under 12-hour days (Fig. 2a) to simulate an estimated threshold for the *sd6* 12-hour flowering causal allele frequency in the non-flowering inbred lines. This simulation predicted that the alternate (non-flowering) allele within *sd6* should be at a high frequency (> 0.8, Fig. S1) in the non-flowering inbred lines. Across all variable sites in the *sd6* interval (6464 total, 5428 genic), 36 variants in twelve genes and 3 SNPs in intergenic regions fit within the modeled threshold and were identified as candidate variants (Fig. 3, Fig. S2). The *FT-*like gene, an *a priori* candidate, has no exonic SNPs differentiating the AHQT and AHQN parents and thus contained no candidate variants. However, one SNP in the gene upstream of *FT* (4.7KB upstream of the *FT* start site, Fig. 3, Fig. S2), has an alternate allele frequency of 0.78, just shy of the modeled threshold for candidates. None of the population genomic candidate variants were within compelling *a priori* functional candidate genes (Fig. 3; Table 4), and the 3 candidate SNPs in intergenic regions were 40KB (1 variant) or 228KB (2 variants) downstream of any functional candidate genes (Fig. S2). Of course, regulatory variants *in cis* with the *sd6* genes, or trans-acting factors not annotated as genes (e.g. small RNAs) which were not well-covered enough to call, may be causal.

The breadth and relative weakness of the *sd11* QTL (Fig. 2) precluded a similar approach to refining candidate variants across the entire region. However, the dual clusters of candidate *MAF*-like genes (MgAHQT1.11G071900 - 72400, MgAHQT1.11G073100-73600) on Chr 11 of the AHQT reference genome exhibited copy number variation when compared to the non-flowering AHQN parental line. Specifically, MgAHQT1.11G072000 appears duplicated in AHQN (average median-standardized coverage = 2.37 at AHQN, 32% of sites heterozygous), while the last two *MAFs* (MgAHQT1.11G073500, MgAHQT1.11G073600) have no exonic coverage in AHQN and are putatively deleted (Fig. S3).

### Fine-scale geothermal gradient predicts 12-hour flowering frequency and phenology in the field

Plants sampled as seeds from AHQT and AHQN extremes and the intervening thermal population (AHQCanyon) at Agrostis Headquarters (AHQ) (Fig. 4a), exhibited significantly different probabilities of 12-hour flowering in growth chambers (p < 0.0001 for all comparisons). A 12-hour minimum daylength for flowering was nearly fixed at AHQT (0.98, 95% CI 0.89-1), common at AHQCanyon (0.73, 95% CI 0.68-0.75), and rare at AHQN (0.11, 95% CI 0-0.31). 85% of AHQT maternal families were entirely flowering (N = 20, n = 1 - 8 per family) and 86% of AHQN maternal families were entirely non-flowering (N = 10, n = 1 - 5 per family) under 12-hour days, suggesting that the causal *sd6* variant is generally homozygous for alternative alleles in each population. In contrast, the intermediate flowering of AHQCanyon families likely reflects a high incidence of wild heterozygotes (as opposed to a mosaic of AHQT and AHQN-like patches), as 98% of maternal families (N = 100, n = 1 – 12 per family) were mixed in their flowering (Fig. 4a).

Like AHQT, the intervening AHQCanyon quadrats are annual due to high soil and air temperatures and severe drought after the end of snowfall. However, at the quadrat scale, both field phenology and environmental data suggest greater variation in the timing and selective effects of geothermal heat and dry-down in the AHQCanyon thermal area (Fig. 3b). Specifically, plants in all AHQT quadrats surveyed (1, 4, 5) and the five upper AHQCanyon quadrats (6-10) reached peak flowering by mid to late April (April 24, 2010; April 15, 2011) and were completely dead by July 9 (Fig 4b). In contrast, the two AHQCanyon quadrats closest to AHQN (12 and 13) reached peak flowering six weeks later than the other thermal sites in 2010 (start May 9, peak June 16). Across AHQT, AHQCanyon, and AHQN, maximum daily plant temperature significantly predicted flowering state (pre-flowering, flowering, or post-flowering) in an ordinal logistic regression (p = 0.003). On June 16, 2010, when flowering was at its peak at the late-flowering AHQCanyon quadrats, 4-PM soil temperature in the two late-flowering quadrats was 7.1 degrees cooler than soil at AHQT and upper AHQCanyon (Fig. 4b). The AHQCanyon plants screened for minimum critical photoperiod were drawn from a continuously sampled transect vs. the fixed environmental quadrats, which precludes direct comparisons of local frequencies and environments. However, fine-scale temperature and moisture variation, even at this very local scale (Fig. 4) appears to allow later flowering at less extreme AHQCanyon quadrats, likely shifting the frequency of sd6 and increasing opportunities for gene flow with later-flowering AHQN.

## Discussion

Flowering plants integrate cues from their environment to initiate flowering (Andrés & Coupland, 2012), and differences in response to environmental cues fine-tune flowering to a plant’s habitat. Shifts in flowering phenology are a ubiquitous response to anthropogenic climate change (Inouye, 2022), but for the most part it is unclear whether these rapid flowering time shifts are plastic or genetic (but see Franks *et al*., 2007). Geothermal soil mosaics provide a natural experiment in plant responses to temperature variation (O’Gorman *et al*., 2014; Leblans *et al*., 2017) without the confounding effects of population structure or ecological and environmental variables that covary with temperature over larger spatial scales. Here, we found that Yellowstone’s mosaic of geothermal habitats generates extensive variation in the key phenological trait of critical daylength for flowering over narrow spatial scales. The most extreme shift to 12-hour flowering is shaped by two novel major loci, both of which contain compelling *a priori* candidate genes for involvement in flowering time. This suggests that although the loci shaping daylength requirements YNP *M. guttatus* are distinct from those underlying other transitions in the species complex, the molecular basis of flowering phenology variation is likely conserved at the molecular level. Furthermore, fine-scale variation in temperature predicts flowering phenology in the field. Together, these analyses demonstrate that fine-scale selection by edaphic (soil) mosaics can generate major shifts in genetically-based flowering time cues in the absence of climatic variation.

### Geothermal inputs shape genetic differences in daylength requirement

Throughout the *M. guttatus* complex, annuals tend to flower early before summer drought (Troth *et al*., 2018), a strategy often facilitated by flowering under shorter days (Friedman & Willis, 2013; Fishman *et al*., 2014). Similarly, Yellowstone thermal annuals generally require shorter minimum critical daylengths than nonthermal perennials (Fig. 1), enabling them to reproduce in spring before the thermal soils become lethally hot and dry. Thus, genetic variation in daylength requirements reflects a match to the novel growing season created by spring snowmelt and summer dry-down near Yellowstone’s geothermal features vs. the snow-enforced summer flowering season of nonthermal perennial populations. Such flowering cue adaptation is common across longitudinal clines (Samis *et al*., 2012; Friedman & Willis, 2013; Ågren *et al*., 2017; Fournier-Level *et al*., 2022) or between co-occurring sister species using distinct microhabitats (Fishman *et al*., 2014; Spriggs *et al*., 2019). Furthermore, Yellowstone *M. guttatus’* habitat-associated MCD variation aligns with widespread prior findings that edaphic mosaics can shape local adaptation in fundamental plant traits (Antonovics, 2006; Rajakaruna, 2018), including flowering time (Sianta & Kay, 2021). However, compared to the abundant edaphic adaptation literature (reviewed in Rajakaruna, 2018), relatively little is known about plant adaptation to geothermal soil mosaics. In Iceland, Geothermal temperatures predict flowering phenology in grasslands (Leblans *et al*., 2017) and in *Ranunculus acris* (Perron, 2017), but *Thymus praecox spp. arcticus* has inconsistent phenological responses to soil temperature, and decreased fitness on hotter soils (Perron, 2017). It is unknown whether the geothermally-associated phenology differences in grasses and *R. acris* are shaped by plasticity, or genetic changes in MCD or other cue responses (Leblans *et al*., 2017; Perron, 2017). In *Cerastium fontanum*, field phenology differences between hotter and cooler sites on a geothermal mosaic are reversed in a common garden; there is evidence for counter-gradient natural selection for later flowering at hotter sites and earlier flowering at cooler sites (Valdés *et al*., 2019; Ehrlén *et al*., 2023). This contrasts with our finding of genetic differences in MCD among YNP thermal annuals and nonthermal perennials, and illustrates that historical contingencies and genetic background shape the evolutionary path of plant populations in geothermal environments.

Edaphic soil mosaics can shape adaptation in the face of gene flow, leading to genetic differentiation at discrete loci and low overall genomic divergence (e.g., Lee & Coop, 2017) However, soil mosaics can also shape divergence in reproductive traits (e.g., Christie & Macnair, 1987; Wright *et al*., 2013), which can ultimately lead to speciation (reviewed in Baack *et al*., 2015). For the most part, we find evidence of edaphic adaptation in the face of gene flow in Yellowstone. Thermal annuals tend to flower under shorter daylengths, but the association is not perfect (some thermal annual lines had 14- or 15-hour daylength requirements, Fig. 1); consistent with the maintenance of high gene flow between neighboring thermal and nonthermal populations (Kolis *et al*., 2022). However, our finding that only AHQT lines flowers under 12-hour days (Fig 1, Fig 4a) indicates that AHQT is distinct from other YNP *M. guttatus*. This shorter daylength requirement, in conjunction with AHQT’s ∼200m geographic isolation from all other *M. guttatus* habitat (Lekberg *et al*., 2012) likely contributes to AHQT’s strong phenotypic and genetic divergence, and may facilitate incipient speciation.

### Daylength requirements are shaped by novel QTLs in Yellowstone *M. guttatus*, but both loci contain highly conserved flowering time genes

Mapping the genetic basis of repeated adaptation, such as shifts in critical daylength for flowering, allows us to address fundamental questions about the repeatability and predictability of evolution (Langerhans & DeWitt, 2004; Orr, 2005). Parallel phenotypic changes can be shaped by the same mutation at the same loci, different mutations at the same loci, or different mutations in distinct and independent loci (Elmer & Meyer, 2011). Re-use of the same loci for parallel phenotypic shifts is more common in closely related species, although exceptions are common (Conte *et al*., 2012; Bohutínská *et al*., 2021). Spatial and temporal variation in selective environment and historical contingencies such as drift and gene flow can lead to distinct genetic architectures or genes shaping repeated evolutionary transitions, even within a species (Kaeuffer *et al*., 2012; Bailey *et al*., 2015; Bohutínská *et al*., 2021). However, repeated association of the same or similar genes shaping traits such as color variation (MC1R genes, Hoekstra & Price, 2004), eye development (opsin genes, Goldsmith, 1990), and flowering time regulation (FT genes, Zeevaart, 2006) suggests that certain traits are under constraint at the molecular pathway level (Storz, 2016). Trait transitions may be governed by distinct and independent loci but still exhibit molecular parallelism, if the loci include the same genes or gene families as those involved in widespread trait transitions.

Two loci (*sd6*, *sd11*) that generate a 2 month shift in minimum critical photoperiod (15 to 12 hours; May 23 to March 21; NOAA, 2025) between adjacent thermal annual (AHQT) and nonthermal perennial (AHQN) monkeyflower populations. These are distinct from three QTLs previously mapped in comparable inter-ecotype and interspecific transitions in yellow monkeyflowers (*sd8a*, *sd8b*; Friedman, 2014, *sd7*, *sd8b*; Fishman *et al*., 2014, respectively). In each of the three studies (which involve crosses between non-overlapping parents), only two major QTLs were sufficient to generate the entire shift (Fig. 3), suggesting a consistent oligogenic architecture for this binary trait. Similarly, all three studies found additivity among QTLs, with non-parental genotypic combinations behaving discretely as individuals (i.e., 12-hr flowering vs. long-term inhibited) but with predictably intermediate probabilities as categories. This suggests a model in which a few major loci act as toggle switches that dramatically raise or lower a barrier (or increase flux over a fixed barrier) to the flowering transition, but smaller environmental and/or genetic factors may be important when both flux and threshold are intermediate. Such simplicity is consistent with the general florigen-as-central-regulator model for flowering (Corbesier *et al*., 2007) and implies that very large differences in the seasonal timing of flowering within and among wild populations can arise from a very few adaptive genetic changes. Similar patterns are seen in *Arabidopsis thaliana* (Putterill *et al*., 2004), *Brassica napus* and *Brassica rapa* (Osborn *et al*., 1997; Cai *et al*., 2008), and rice (Yano *et al*., 2001) where wide variation in flowering time is under the control of only a few loci.

Long-term balancing selection and introgression are increasingly evident as important contributors to repeated adaptation (Blount *et al*., 2018) even over large spatial and temporal scales (Mäkinen *et al*., 2008; Delph & Kelly, 2014; Jones *et al*., 2018); these processes could have led to the sharing of causal loci between YNP *M. guttatus* and previously mapped transitions. However, our major 12-hour flowering QTLs do not overlap with parallel transitions in the *M. guttatus* complex, suggesting that the geographical isolation of Yellowstone may have prevented sharing of existing alleles (e.g., from nonthermal annuals in highly seasonal habitats in California). Alternatively, the ecological uniqueness of Yellowstone’s thermal annual habitats may have precluded the spread of more 12-hour flowering widespread variants even if available via gene flow. Furthermore, the known *M. guttatus* flowering time loci on chromosomes 7 and 8 may have pleiotropic effects on other traits (or are in tight linkage with loci affecting multiple traits), which could result in decreased fitness in Yellowstone’s harsh and unusual environment, and favor the use of a distinct locus for flowering time.

The lack of allele- or locus-sharing with parallel shifts elsewhere in the *M. guttatus* complex (Fishman *et al*., 2014; Friedman, 2014) suggests that a relatively large portfolio of loci may contribute to simple evolutionary jumps to 12-hour flowering. However, at broader trait and pathway levels, there may be greater predictability. The *sd6* and *sd11* regions have not been associated specifically with 12-hour flowering in other *M. guttatus* complex work, but *sd6* is associated with days to flower under long days (Friedman *et al*., 2015), and *sd11* overlaps with a QTL for photoperiod-dependent vernalization, which interacts epistatically with photoperiod loci (Friedman & Willis, 2013). This suggests that distinct variants in the same underlying gene(s) may influence other aspects of flowering phenology, implying a finite set of *M. guttatus* loci as targets for adaptive mutations in flowering time range-wide. Furthermore, both QTLs contain strong functional candidates for modulation of flowering (*sd6* = FT-like, *sd11* = MAF cluster), suggesting broader molecular constraints on environmental flowering cues (Srikanth & Schmid, 2011; Pin & Nilsson, 2012; Putterill & Varkonyi-Gasic, 2016).

Although finer-mapping and genetic manipulation will be necessary to pin down the causal variation underlying our major QTLs, each contains a strong candidate gene family with known flowering time functions. In diverse angiosperms, *FT*-like genes encode the florigen, a leaf-to-meristem transmissible flowering signal (summarized in Zeevaart, 2006) whose concentration integrates signals from multiple internal and environmental inputs to trigger flowering (Pin & Nilsson, 2012; Putterill & Varkonyi-Gasic, 2016). In addition to encoding florigen itself, paralogous *FT*-like genes can act as both promoters and repressors of flowering (Yamaguchi *et al*., 2005; Pin *et al*., 2010; Putterill & Varkonyi-Gasic, 2016), so are an evolutionarily labile target for selection on flowering time. *FT-like* genes have been associated with flowering time in nearly every angiosperm species studied (Kojima *et al*., 2002; Ratcliffe *et al*., 2003; Lifschitz & Eshed, 2006; Laurie *et al*., 2011; Pin & Nilsson, 2012; Putterill & Varkonyi-Gasic, 2016; Zhou *et al*., 2019; Liang *et al*., 2019; Ning *et al*., 2019; Zhao *et al*., 2022), including 12-hour flowering in *M. nasutus* x *M. guttatus* hybrids (Fishman *et al*., 2014). The *FT*-like gene under the sd6 peak has no coding variants between the AHQT and AHQN QTL-mapping parents (Fig. 3) and no variants in the inbred line set fitting modeled allele frequencies for non-flowering lines. However, *FT* expression is often shaped by variation in its cis-regulatory regions (Liu *et al*., 2014; Luo *et al*., 2019) and the abundant variation in flanking intergenic regions may be causal. The *sd11* candidate genes, a cluster of *MADS AFFECTING FLOWERING* transcription factors (MAFs) exhibit more obvious candidate variants: copy number variation (CNV) between AHQT and AHQN at three of the 10 genes in the chr11 *MAF* cluster (Fig. S3). MAFs affect flowering through regulation *FT* in many systems (Ratcliffe *et al*., 2003; Yan *et al*., 2004; Deng *et al*., 2011; Theißen *et al*., 2016; Liang *et al*., 2019), acting as both repressors and promoters (Andrés & Coupland, 2012). Copy number variants (CNVs) often contribute to adaptation to novel environments (Iskow *et al*., 2012; Hull *et al*., 2017) and CNVs in *MAF*s are known to shape vernalization and flowering time in *Arabis* (Madrid *et al*., 2021). As additional reference genomes and transcriptomic datasets are developed for diverse yellow monkeyflowers (e.g., (Lovell *et al.,* 2025), it will become increasingly possible to uncover the causal variants underlying *sd6* and s*d11* in Yellowstone, as well as understand the flowering consequences of copy number variation in *MAF* and *FT* families in *M. guttatus* complex range-wide.

### Fine-scale temperature variation shapes flowering phenology on the landscape

Yellowstone’s geothermal soils provide the rare opportunity to observe adaptive divergence in flowering time in its incipient stages, identify the environmental factors exerting divergent selection, and quantify the degree of within-population segregation for flowering time traits. Here, we discovered that plant temperature significantly predicts flowering phenology at the meter scale in AHQ (Fig. 4b), and the frequency of 12-hour flowering broadly mirrors this gradient in temperature and moisture from the highly thermal AHQT to the nonthermal bog AHQN (Fig. 4a). Interestingly, most maternal families in AHQCanyon are segregating for flowering under 12-hour days, suggesting an abundance of *sd6* and/or *sd11* heterozygotes. Both the patchy distribution of cooler and warmer thermal sites in AHQCanyon (Fig. 4b) and its proximity to nonthermal habitat may contribute to the non-fixation of 12-hour flowering in this thermal annual site. Overall, YNP *M. guttatus* populations exhibit remarkable evolutionary lability for daylength requirements, tightly matching flowering phenology to environmental conditions.

## Conclusions

The diversity of *M. guttatus’* daylength requirements for flowering across Yellowstone microgeographic soil mosaic illustrates that this key life history trait readily responds to selection, even with high gene flow. Just as continent-scale environmental variation has shaped clines in flowering phenology across species ranges (Stinchcombe *et al*., 2004; Samis *et al*., 2012; Friedman & Willis, 2013), the interaction between soil temperature and moisture exerts variable selection on daylength requirements throughout Yellowstone on very short spatial scales. Novel 12-hour flowering QTLs (vs. previous studies in *M. guttatus* complex) broaden the set of *Mimulus* genomic targets for shifts to extreme spring flowering, but overlap with QTLs for other phenological traits and strong functional candidates suggest conservation at the pathway vs. locus level. Although additional work will be necessary to fully connect the underlying molecular variants with environmentally-dependent flowering phenotypes and fitness, these experiments illustrate how soil mosaics can select on relatively simple genetic architectures to generate and maintain strong divergence in flowering time cues, even in the absence of climate variation and in the face of ongoing gene flow.

## Acknowledgements

The authors gratefully acknowledge Thom Nelson for his advice on the molecular and analytical work and Kailey Baisen for her assistance in the greenhouse. Funding support was provided by NSF grants NSF DEB-1457763 and by NSF OIA-1736249 to Zac Cheviron, Lila Fishman, Jay Storz, Kristi Montooth, and Jeffrey Good. Field collections were conducted under Yellowstone National Park Staff for research permits (YELL-2010 - 2018-SCI-5834). The AHQTv1 reference genome assembly and annotation (proposal: 10.46936/10.25585/60001364) was conducted by the U.S. Department of Energy Joint Genome Institute (https://ror.org/04xm1d337), a DOE Office of Science User Facility, is supported by the Office of Science of the U.S. Department of Energy operated under Contract No. DE-AC02-05CH11231.

## Author Contributions

CSB—data collection, analysis, visualization, drafting of manuscript. MH—data collection, KK— data collection, KS—data collection, FF—data collection, LF—conceptualization, analysis, interpretation, drafting and editing of manuscript.

## Data availability

The raw sequence data (PRJNA1050826, PRJNA1051082) are publicly available. Datasets for all analyses are archived at Dryad and will be fully released upon publication.

---

**Supplementary Figure 1.**
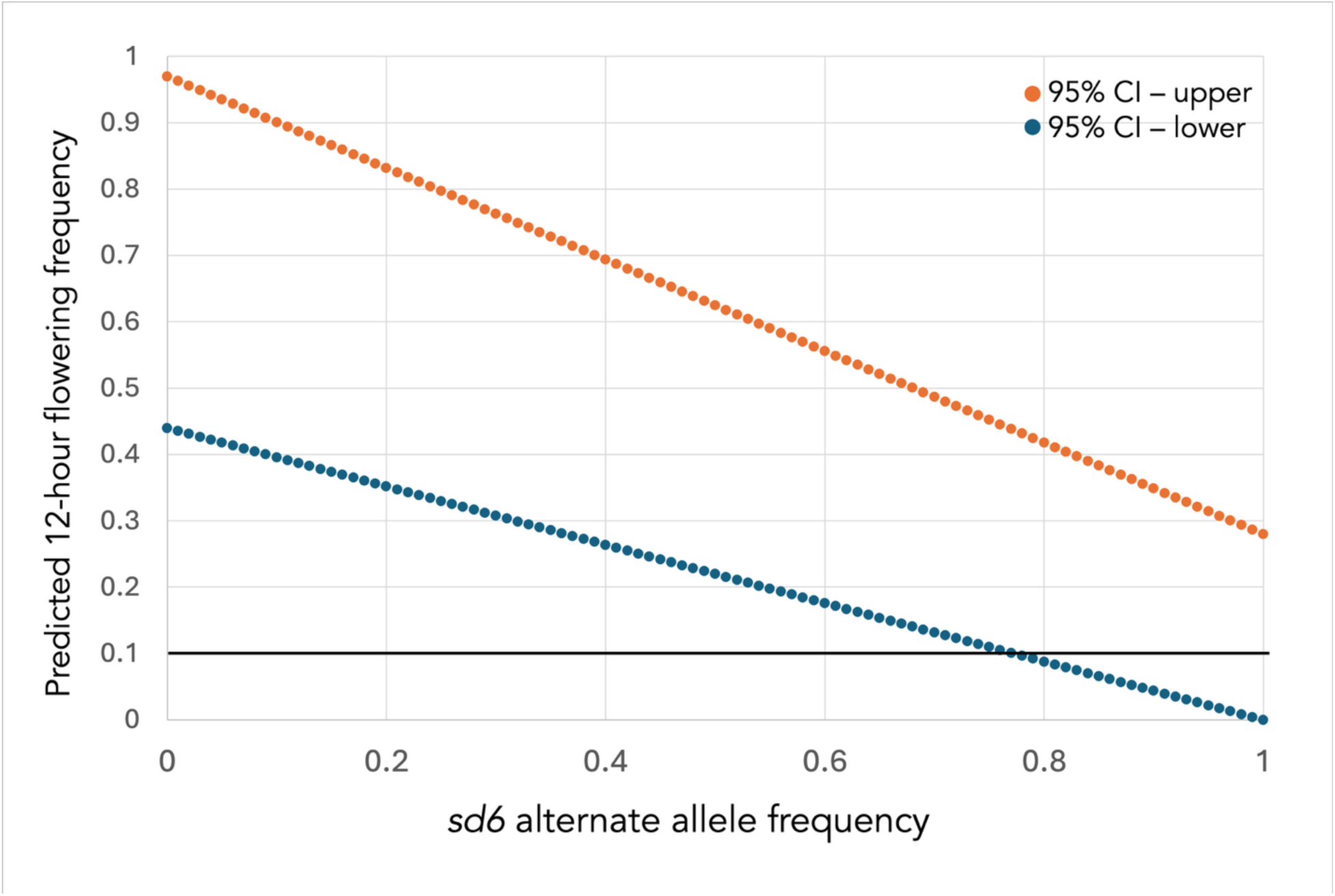
Simulation of the frequency of causal allele within *sd6* in non-flowering inbred lines. Predicted frequency of the alternate (non-flowering) allele in the 44 inbred lines which did not flower under 12-hour days. Conservatively assuming that the inbred lines are fixed for the non-flowering allele at *sd11*, we calculated the predicted flowering frequency at alternate allele frequencies ranging from 0 – 1, using the 95% confidence intervals for the frequency at which TT and NN individuals flower under 12-hour days from the QTL map (Fig. 2b, TT flowering = 0.44 – 0.97; NN flowering = 0 – 0.28). Because the none of the 44 inbred lines flowered under 12-hour days (95% CI = 0 – 0.1), the alternate allele frequency for the causal variant should be > 0.8; this threshold is marked with a horizontal line.

**Supplementary Figure 2.**
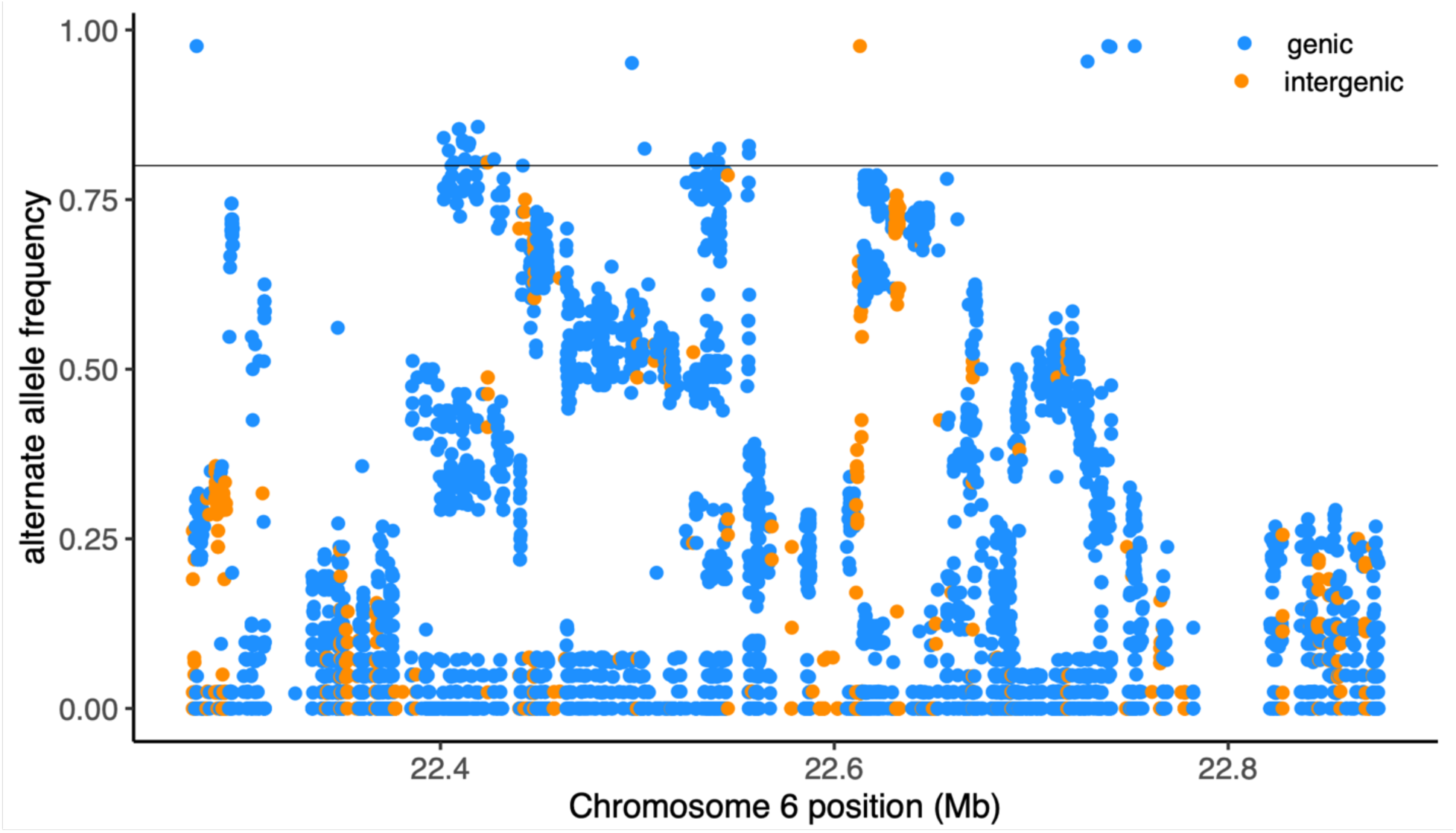
Alternate allele frequency in within *sd6* in non-flowering inbred lines. Frequency of the alternate (non-flowering) allele for each variant within the *sd6* QTL in non-12-hour-flowering Yellowstone *Mimulus guttatus*. Candidate variants are those with an alternate allele frequency ≥ 0.8.

**Supplementary Figure 3.**
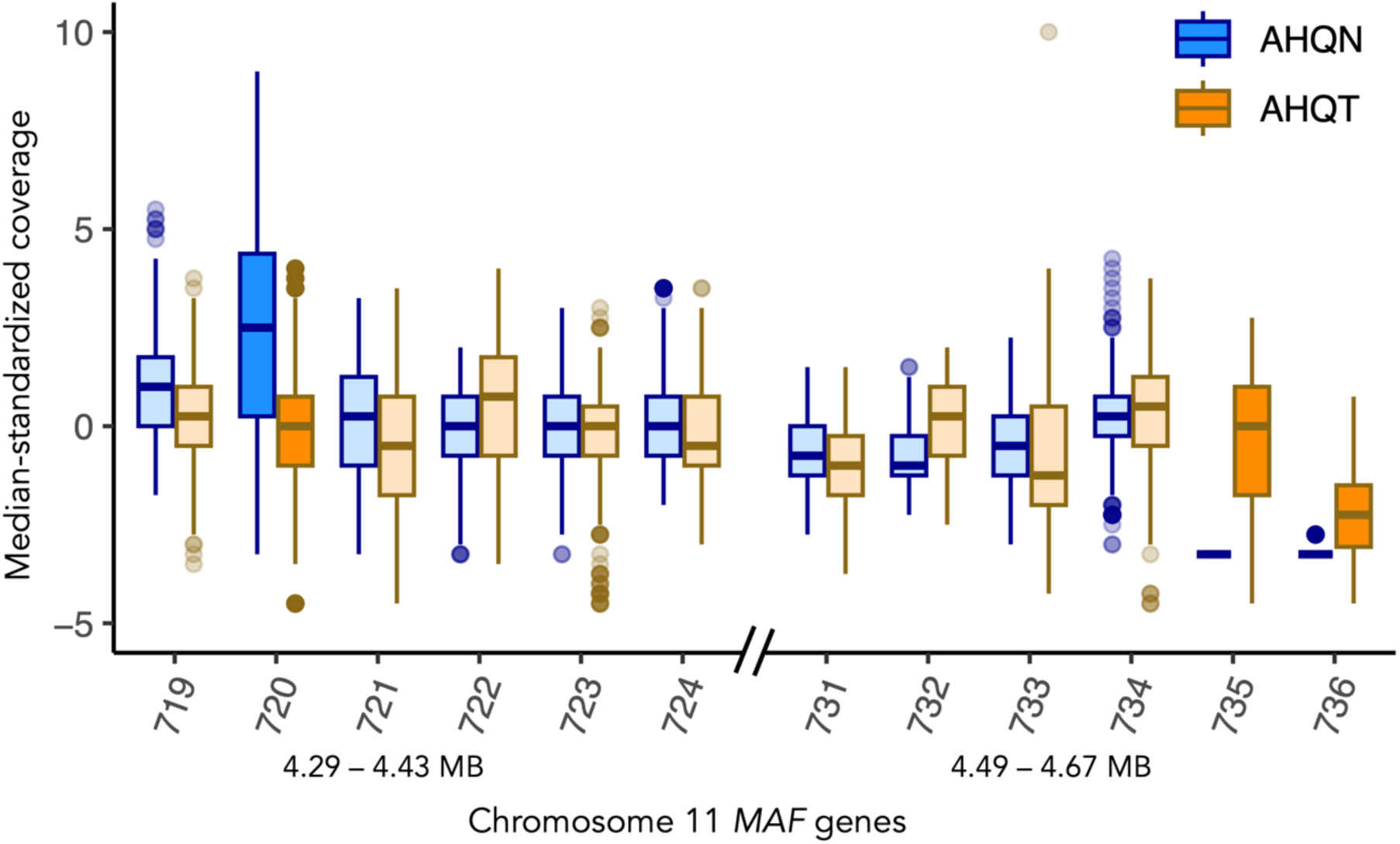
Coverage differences indicate copy number variants between thermal and nonthermal lines in a *MAF* gene cluster. Median-standardized coverage in AHQT and AHQN whole genome sequenced inbred lines in the two tandem clusters of *MADS AFFECTING FLOWERING* transcription factors within the *sd11* QTL. Boxplots with darker colors are genes with putative copy number variants or presence-absence variants between AHQT and AHQN: MgAHQTv1.11G72000 appears duplicated in AHQN relative to AHQT, and MgAHQTv1.11G73500 and MgAHQTv1.11G73600 are present in AHQT but absent in AHQN.

**Supplementary Table 1.** Inbred lines, their habitat of origin, (assigned by and the analyses in which each line is included. SRA accession numbers are provided for whole-genome sequenced lines.

| Line Name | Habitat | Latitude | Longitude | Screening | Accession |
| --- | --- | --- | --- | --- | --- |
| AHQN | Nonthermal Perennial | 44.435226 | -110.8092 | 12-15h | SRR33733352 |
| AHQT | Thermal Annual | 44.431354 | -110.81474 | 12h | SRR486613 |
| YGBR_10.1 | Nonthermal Perennial | 44.70995 | -110.7412 | 12-13h | SRR27166891 |
| YGBR_12.1 | Nonthermal Perennial | 44.65036 | -110.8458 | 12-15h | SRR27166863 |
| YGBR_3.1 | Nonthermal Perennial | 44.68078 | -110.7445 | 12-15h | SRR27166841 |
| YGBR_4.3 | Nonthermal Perennial | 44.68063 | -110.7446 | 12-15h | SRR27166830 |
| YGBR_7.1 | Nonthermal Perennial | 44.68967 | -110.7456 | 12-15h | SRR27166819 |
| YHLB_11.1 | Nonthermal Perennial | 44.29819 | -110.5172 | 12h | SRR27166890 |
| YHLB_13.3 | Nonthermal Perennial | 44.30096 | -110.51966 | 12h | o |
| YHLB_14.5 | Thermal Annual | 44.30876 | -110.526 | 12-15h | o |
| YHLB_15.1 | Thermal Annual | 44.3134 | -110.5361 | 12-15h | SRR27166871 |
| YHLB_2.1 | Nonthermal Perennial | 44.28149 | -110.5065 | 12-15h | SRR27166870 |
| YHLB_3.4 | Thermal Annual | 44.28164 | -110.507 | 12h | SRR27166869 |
| YHLB_5.5 | Nonthermal Perennial | 44.28288 | -110.50556 | 12-15h | o |
| YHLB_6.2 | Nonthermal Perennial | 44.28996 | -110.50391 | 14-15h | o |
| YLGB_1.1 | Thermal Annual | 44.53564 | -110.7998 | 12-15h | SRR27166866 |
| YLGB_18.2 | Nonthermal Perennial | 44.55012 | -110.8065 | 12h | SRR27166864 |
| YLGB_24.3 | Nonthermal Perennial | 44.5335 | -110.8072 | 12h | SRR27166857 |
| YLGB_25.1 | Thermal Annual | 44.53369 | -110.8069 | 12-15h | SRR27166856 |
| YLGB_31.1 | Nonthermal Perennial | 44.53288 | -110.79717 | 12-15h | SRR27166851 |
| YLGB_4.2 | Thermal Annual | 44.53504 | -110.8009 | 12h | SRR27166850 |
| YLSG_2.1 | Thermal Annual | 44.41638 | -110.8091 | 12-15h | SRR27166847 |
| YLSG_4.4 | Thermal Annual | 44.4167 | -110.8134 | 12-15h | SRR27166846 |
| YLSG_5.4 | Nonthermal Perennial | 44.4164 | -110.8138 | 12-15h | SRR27166845 |
| YMGB_13.3 | Thermal Annual | 44.52787 | -110.8366 | 12-15h | SRR27166842 |
| YMGB_6.1 | Thermal Annual | 44.51635 | -110.833 | 12-15h | SRR27166840 |
| YOJO_1.5 | Thermal Annual | 44.5629 | -110.8387 | 12-15h | SRR27166839 |
| YOJO_2.2 | Thermal Annual | 44.56269 | -110.8389 | 12-15h | SRR27166838 |
| YOJO_4.1 | Thermal Annual | 44.56162 | -110.8358 | 12-15h | SRR27166836 |
| YOJO_5.1 | Thermal Annual | 44.5613 | -110.8353 | 12-15h | SRR27166835 |
| YOJO_6.3 | Thermal Annual | 44.56116 | -110.8349 | 12-15h | o |
| YOJO_7.1 | Nonthermal<br>Perennial | 44.56063 | -110.8346 | 12-15h | SRR27166834 |
| YSGB_2.2 | Thermal Annual | 44.35524 | -110.7978 | 12-13h | SRR27166831 |
| YSGB_4.4 | Thermal Annual | 44.35484 | -110.7991 | 12-15h | o |
| YSGB_5.1 | Nonthermal<br>Perennial | 44.35411 | -110.7985 | 12-15h | SRR27166829 |
| YSGB_8.2 | Nonthermal<br>Perennial | 44.35206 | -110.8007 | 14-15h | o |
| YSGB_9.4 | Thermal Annual | 44.35172 | -110.8001 | 12-15h | SRR27166826 |
| YUGB_10.1 | Thermal Annual | 44.46513 | -110.837 | 12-15h | SRR27166825 |
| YUGB_13.2 | Thermal Annual | 44.46898 | -110.8406 | 12-15h | SRR27166824 |
| YUGB_15.1 | Thermal Annual | 44.47285 | -110.8415 | 12-13h | SRR27166823 |
| YUGB_19.2 | Nonthermal<br>Perennial | 44.47575 | -110.8443 | 12-15h | o |
| YUGB_20.1 | Thermal Annual | 44.47925 | -110.8489 | 12-13h | SRR27166818 |
| YUGB_23.2 | Nonthermal<br>Perennial | 44.47966 | -110.8489 | 12-15h | SRR27166815 |
| YUGB_24.1 | Nonthermal<br>Perennial | 44.46165 | -110.8543 | 14-15h | o |
| YUGB_25.2 | Thermal Annual | 44.46138 | -110.855 | 12-15h | SRR27166813 |
| YUGB_8.2 | Thermal Annual | 44.46486 | -110.8371 | 12-15h | SRR27166887 |
| YVTC_2.2 | Nonthermal<br>Perennial | 44.65226 | -110.5452 | 12-15h | SRR27166886 |
| YVTC_3.2 | Thermal Annual | 44.65064 | -110.5577 | 12-15h | SRR27166885 |
| YVTC_6.1 | Thermal Annual | 44.65031 | -110.5569 | 12-15h | SRR27166884 |
| YVTC_7.2 | Thermal Annual | 44.65005 | -110.5564 | 12-15h | SRR27166883 |
| YVTC_8.2 | Thermal Annual | 44.65033 | -110.55583 | 12-15h | o |
| YWTB_2.2 | Thermal Annual | 44.41591 | -110.5697 | 12-15h | SRR27166879 |
| YWTB_3.2 | Nonthermal<br>Perennial | 44.41591 | -110.5697 | 12-15h | SRR27166877 |
| YWTB_4.2 | Thermal Annual | 44.41825 | -110.5721 | 12-15h | o |
| YWTB_6.2 | Nonthermal<br>Perennial | 44.41709 | -110.5732 | 13h | o |
| YWTB_7.2 | Nonthermal<br>Perennial | 44.41642 | -110.57293 | 12-15h | SRR27166874 |
| YWTB_8.2 | Thermal Annual | 44.41563 | -110.5724 | 12-15h | SRR27166873 |

**Supplementary Table 2.** *sd6* gene annotations. Names, base pair coordinates and annotations of all genes within the *sd6* QTL, using terms from the AHQTv1 reference genome.

| Gene | Chr | bp_start | bp_end | function |
| --- | --- | --- | --- | --- |
| MgAHQT1.06G197500 | Chr_06 | 22270071 | 22271597 | Protein of unknown function |
| MgAHQT1.06G197600 | Chr_06 | 22275271 | 22280934 | mitochondrial substraits carrier family protein |
| MgAHQT1.06G197700 | Chr_06 | 22282689 | 22283810 | F-box and associated interaction domains-containing protein |
| MgAHQT1.06G197800 | Chr_06 | 22288033 | 22289190 | F-box and associated interaction domains-containing protein |
| MgAHQT1.06G197900 | Chr_06 | 22292000 | 22295357 | Protein IQ-domain 3 related |
| MgAHQT1.06G198000 | Chr_06 | 22297665 | 22302692 | MYB family transcription factor |
| MgAHQT1.06G198100 | Chr_06 | 22303640 | 22308500 | SET domain protein |
| MgAHQT1.06G198200 | Chr_06 | 22309912 | 22311314 | cystein-rich repeat secretory protein 11-related |
| MgAHQT1.06G198300 | Chr_06 | 22316586 | 22324675 | leucine-rich repeat (LRR) family protein |
| MgAHQT1.06G198400 | Chr_06 | 22323659 | 22328999 | leucine-rich repeat (LRR) family protein |
| MgAHQT1.06G198500 | Chr_06 | 22328978 | 22333207 | NB-ARC domain leucine-rich repeat |
| MgAHQT1.06G198600 | Chr_06 | 22334763 | 22341711 | AAA ATPase domain |
| MgAHQT1.06G198700 | Chr_06 | 22343009 | 22349027 | leucine-rich repeat (LRR) family protein |
| MgAHQT1.06G198800 | Chr_06 | 22349812 | 22351303 | leucine-rich repeat (LRR) family protein |
| MgAHQT1.06G198900 | Chr_06 | 22358568 | 22361551 | leucine-rich repeat (LRR) family protein |
| MgAHQT1.06G199000 | Chr_06 | 22362829 | 22365971 | leucine-rich repeat (LRR) family protein |
| MgAHQT1.06G199100 | Chr_06 | 22368127 | 22372734 | leucine-rich repeat (LRR) family protein |
| MgAHQT1.06G199200 | Chr_06 | 22372672 | 22376835 | small nuclear ribonuclear protein |
| MgAHQT1.06G199300 | Chr_06 | 22384352 | 22386992 | GDSL esterase / lipase LDL1 |
| MgAHQT1.06G199400 | Chr_06 | 22387846 | 22390873 | zinc finger FYVE domain containing protein |
| MgAHQT1.06G199500 | Chr_06 | 22392236 | 22398935 | plastid transcriptionally active 3 |
| MgAHQT1.06G199600 | Chr_06 | 22399238 | 22422948 | Translational activator GCN1 |
| MgAHQT1.06G199700 | Chr_06 | 22424873 | 22429432 | Protein of unknown function |
| MgAHQT1.06G199800 | Chr_06 | 22430153 | 22432394 | exosome complex component CSL4 |
| MgAHQT1.06G199900 | Chr_06 | 22432611 | 22435217 | exosome complex component CSL4 |
| MgAHQT1.06G200000 | Chr_06 | 22440148 | 22442373 | Ammonium transporter |
| MgAHQT1.06G200100 | Chr_06 | 22445039 | 22447513 | Ammonium transporter |
| MgAHQT1.06G200200 | Chr_06 | 22447756 | 22449542 | family not named |
| MgAHQT1.06G200300 | Chr_06 | 22450430 | 22455138 | Aminopyrimidine aminohydrolase / thiaminase II |
| MgAHQT1.06G200400 | Chr_06 | 22463442 | 22467648 | leucine-rich repeat receptor-like protein kinase PXL1 |
| MgAHQT1.06G200500 | Chr_06 | 22469063 | 22481949 | transcriptional regulator ATRX |
| MgAHQT1.06G200600 | Chr_06 | 22481903 | 22484789 | Protein of unknown function |
| MgAHQT1.06G200700 | Chr_06 | 22486350 | 22488063 | calcium-dependent lipid-binding domain-containing protein-related |
| MgAHQT1.06G200800 | Chr_06 | 22489275 | 22490900 | prostaglandin-endoperoxide synthase |
| MgAHQT1.06G200900 | Chr_06 | 22491699 | 22493161 | calcium-dependent lipid-binding domain-containing protein-related |
| MgAHQT1.06G201000 | Chr_06 | 22495556 | 22499430 | disease resistance protein RPM1 |
| MgAHQT1.06G201100 | Chr_06 | 22500623 | 22503866 | leucine-rich repeat (LRR) family protein |
| MgAHQT1.06G201200 | Chr_06 | 22504517 | 22506680 | family not named |
| MgAHQT1.06G201300 | Chr_06 | 22509119 | 22512721 | violaxanthin de-epoxidase (VDE, NPQ1) |
| MgAHQT1.06G201400 | Chr_06 | 22513197 | 22516653 | interleukin-1 receptor associated kinase |
| MgAHQT1.06G201500 | Chr_06 | 22517184 | 22519153 | family not named |
| MgAHQT1.06G201600 | Chr_06 | 22520229 | 22522190 | UDP-arabinopyranose mutase |
| MgAHQT1.06G201700 | Chr_06 | 22523567 | 22526979 | protein kinase family protein |
| MgAHQT1.06G201800 | Chr_06 | 22528484 | 22532970 | cytochrome P450 |
| MgAHQT1.06G201900 | Chr_06 | 22533649 | 22545128 | guanyl-nucleotide exchange factor / SH3 domain-containing protein |
| MgAHQT1.06G202000 | Chr_06 | 22555343 | 22557740 | l-ascorbate oxidase |
| MgAHQT1.06G202100 | Chr_06 | 22558287 | 22562247 | DO-like 15 protein related |
| MgAHQT1.06G202200 | Chr_06 | 22563124 | 22567989 | Six-hairpin glycosidases superfamily protein |
| MgAHQT1.06G202300 | Chr_06 | 22584274 | 22588581 | beta-hexosaminidase |
| MgAHQT1.06G202400 | Chr_06 | 22604971 | 22605423 | probable lipid transfer |
| MgAHQT1.06G202500 | Chr_06 | 22606083 | 22610441 | ATP-dependent RNA helicase |
| MgAHQT1.06G202600 | Chr_06 | 22614159 | 22620926 | LETM1 and EF-hand domain containing protein 1, mitochondrial |
| MgAHQT1.06G202700 | Chr_06 | 22621089 | 22626279 | leucine-rich repeat (LRR) family protein |
| MgAHQT1.06G202800 | Chr_06 | 22626913 | 22630331 | vacuolar ATP synthase subunit H family protein |
| MgAHQT1.06G202900 | Chr_06 | 22633864 | 22636080 | NB_ARC domain |
| MgAHQT1.06G203000 | Chr_06 | 22636271 | 22640035 | Ribosomal protein S5 family protein |
| MgAHQT1.06G203100 | Chr_06 | 22640198 | 22644276 | RNA binding (RRM/RBD/RNP motifs) family protein |
| MgAHQT1.06G203200 | Chr_06 | 22644575 | 22645462 | transmembrane protein |
| MgAHQT1.06G203300 | Chr_06 | 22645485 | 22649477 | Plant stearyl-acyl-carrier-protein desaturase family protein |
| MgAHQT1.06G203400 | Chr_06 | 22651794 | 22653060 | homologous to Flowering locus T gene |
| MgAHQT1.06G203500 | Chr_06 | 22656491 | 22658273 | HXXXD-type acyl-transferase family protein |
| MgAHQT1.06G203600 | Chr_06 | 22660471 | 22666791 | E3 ubiquitin-protein ligase ARI5-related |
| MgAHQT1.06G203700 | Chr_06 | 22666697 | 22670141 | calcineurin-like metallo-phosphoesterase superfamily protein |
| MgAHQT1.06G203800 | Chr_06 | 22670294 | 22672258 | Protein of unknown function |
| MgAHQT1.06G203900 | Chr_06 | 22672316 | 22675793 | AP2-like ethylene-responsive transcription factor |
| MgAHQT1.06G204000 | Chr_06 | 22679252 | 22682598 | phospholipase A(1) LCAT3 |
| MgAHQT1.06G204100 | Chr_06 | 22682849 | 22687028 | TATA box-binding protein-associated factor RNA polymerase subunit B |
| MgAHQT1.06G204200 | Chr_06 | 22687184 | 22689684 | NADH dehydrogenase (ubiquinone) 1 alpha/beta subcomplex 1 |
| MgAHQT1.06G204300 | Chr_06 | 22690271 | 22693807 | calreticulin-3 |
| MgAHQT1.06G204400 | Chr_06 | 22693917 | 22697911 | no definition available |
| MgAHQT1.06G204500 | Chr_06 | 22698427 | 22703689 | LUC7 related protein |
| MgAHQT1.06G204600 | Chr_06 | 22703814 | 22705784 | Protein of unknown function |
| MgAHQT1.06G204700 | Chr_06 | 22708877 | 22709774 | protein TIC 20-II, chloroplastic |
| MgAHQT1.06G204800 | Chr_06 | 22709774 | 22713128 | chloroplastic acetylcoenzyme A carboxylase 1 |
| MgAHQT1.06G204900 | Chr_06 | 22713395 | 22715393 | thioredoxin F2, chloroplast precursor |
| MgAHQT1.06G205000 | Chr_06 | 22714667 | 22718289 | proteinaceous RNase P |
| MgAHQT1.06G205100 | Chr_06 | 22719292 | 22725115 | aspartic proteinase-like protein-2 related |
| MgAHQT1.06G205200 | Chr_06 | 22724886 | 22726797 | Protein of unknown function |
| MgAHQT1.06G205300 | Chr_06 | 22726347 | 22730711 | serine carboxypeptidase-like clase |
| MgAHQT1.06G205400 | Chr_06 | 22730873 | 22732235 | dirigent protein 24-related |
| MgAHQT1.06G205500 | Chr_06 | 22732306 | 22736120 | cytosine deaminase |
| MgAHQT1.06G205600 | Chr_06 | 22737285 | 22741087 | protein phosphatase 2C family protein |
| MgAHQT1.06G205700 | Chr_06 | 22749362 | 22750068 | gamma-glutamylcyclotransferase |
| MgAHQT1.06G205800 | Chr_06 | 22750025 | 22750918 | crib domain-containing protein RIC4 |
| MgAHQT1.06G205900 | Chr_06 | 22751342 | 22753554 | pentatricopeptide repeat domain |
| MgAHQT1.06G206000 | Chr_06 | 22753575 | 22756997 | eIF4-gamma/eIF5/eIF2-epsilon (W2) |
| MgAHQT1.06G206100 | Chr_06 | 22759976 | 22761293 | peroxidase precursor |
| MgAHQT1.06G206200 | Chr_06 | 22761579 | 22763029 | peroxidase precursor |
| MgAHQT1.06G206300 | Chr_06 | 22766788 | 22768391 | peroxidase precursor |
| MgAHQT1.06G206400 | Chr_06 | 22768642 | 22769765 | peroxidase / lactoperoxidase |
| MgAHQT1.06G206500 | Chr_06 | 22778746 | 22780019 | peroxidase / lactoperoxidase |
| MgAHQT1.06G206600 | Chr_06 | 22781576 | 22787910 | peroxidase / lactoperoxidase |
| MgAHQT1.06G206700 | Chr_06 | 22789653 | 22791076 | peroxidase precursor |
| MgAHQT1.06G206800 | Chr_06 | 22818890 | 22820351 | peroxidase precursor |
| MgAHQT1.06G206900 | Chr_06 | 22820829 | 22827552 | coatamer, subunit beta (COPB1, SEC26) |
| MgAHQT1.06G207000 | Chr_06 | 22836348 | 22839311 | WRKY transcription factor 10-related |
| MgAHQT1.06G207100 | Chr_06 | 22840216 | 22841685 | adipose regulatory protein, SEIPIN |
| MgAHQT1.06G207200 | Chr_06 | 22841767 | 22844275 | isopentenyl-diphosphate delta-isomerase |
| MgAHQT1.06G207300 | Chr_06 | 22846672 | 22847973 | zinc finger (C2H2 type) family protein |
| MgAHQT1.06G207400 | Chr_06 | 22851193 | 22855531 | X-box transcription factor related |
| MgAHQT1.06G207500 | Chr_06 | 22856287 | 22856839 | Family not named |
| MgAHQT1.06G207600 | Chr_06 | 22857672 | 22861110 | domain of unknown function |
| MgAHQT1.06G207700 | Chr_06 | 22861322 | 22863566 | Ubiquitin carboxyl-terminal hydrolase family protein |
| MgAHQT1.06G207800 | Chr_06 | 22863584 | 22865732 | ATP hydrolysis coupled transmembrane transport |
| MgAHQT1.06G207900 | Chr_06 | 22867274 | 22868993 | auxin efflux carrier family protein |
| MgAHQT1.06G208000 | Chr_06 | 22873476 | 22876213 | leucine-rich repeat receptor-like protein kinase PXL1 |

**Supplementary Table 3.** Field photographs in quadrats in Yellowstone National Park from the highly thermal habitat AHQT, the thermal AHQCanyon, and nonthermal AHQN in 2010 and 2011.

| Photo | Quad Number | Quad Name | Date | Flower Number | Area | Phenology | Notes |
| --- | --- | --- | --- | --- | --- | --- | --- |
| DSCN4310 | 4 | Lila2 4 | 4/25/10 | 477 | AHQT | early |  |
| DSCN4323 | 5 | Log 5 | 4/25/10 | 398 | AHQT | early |  |
| DSCN4326 | 6 | Canyon 6 | 4/25/10 | 846 | AHQCanyon | early |  |
| DSCN4331 | 7 | Canyon2 7 | 4/25/10 | 447 | AHQCanyon | early |  |
| DSCN4336 | 8 | CFH 8 | 4/25/10 | 661 | AHQCanyon | early |  |
| DSCN4338 | 9 | CFM 9 | 4/25/10 | 703 | AHQCanyon | early |  |
| DSCN4340 | 10 | CFL 10 | 4/25/10 | 69 | AHQCanyon | early |  |
| DSCN4373(1) | 1 | Ylva 1 | 5/8/10 | 6 | AHQT | early |  |
| DSCN4379(1) | 5 | Log 5 | 5/8/10 | 261 | AHQT | early |  |
| DSCN4425 | 12 | Bison 12 | 5/9/10 | 74 | AHQCanyon | late |  |
| DSCN4470(1) | 1 | Ylva 1 | 6/6/10 | 0 | AHQT | early |  |
| DSCN4481(1) | 4 | Lila2 4 | 6/6/10 | 56 | AHQT | early |  |
| DSCN4492(1) | 8 | CFH 8 | 6/6/10 | 75 | AHQCanyon | early |  |
| DSCN4495(1) | 9 | CFM 9 | 6/6/10 | 142 | AHQCanyon | early | overexposed |
| DSCN4656 | 7 | Canyon2 7 | 6/15/10 | 231 | AHQCanyon | early |  |
| DSCN4645 | 1 | Ylva 1 | 6/16/10 | 0 | AHQT | early |  |
| DSCN4649 | 4 | Lila2 4 | 6/16/10 | 31 | AHQT | early |  |
| DSCN4653 | 5 | Log 5 | 6/16/10 | 4 | AHQT | early |  |
| DSCN4655 | 6 | Canyon 6 | 6/16/10 | 29 | AHQCanyon | early |  |
| DSCN4657 | 8 | CFH 8 | 6/16/10 | 58 | AHQCanyon | early |  |
| DSCN4658 | 9 | CFM 9 | 6/16/10 | 112 | AHQCanyon | early |  |
| DSCN4659 | 10 | CFL 10 | 6/16/10 | 0 | AHQCanyon | early |  |
| DSCN4662(1) | 13 | Boulder 13 | 6/16/10 | 613 | AHQCanyon | late |  |
| DSCN4665(1) | 12 | Bison 12 | 6/16/10 | 419 | AHQCanyon | late |  |
| DSCN4730 | 9 | CFM 9 | 7/9/10 | 0 | AHQCanyon | early |  |
| DSCN4731 | 10 | CFL 10 | 7/9/10 | 0 | AHQCanyon | early |  |
| DSCN4736 | 13 | Boulder 13 | 7/9/10 | 0 | AHQCanyon | late |  |
| DSCN4740 | 12 | Bison 12 | 7/9/10 | 0 | AHQCanyon | late |  |
| DSCN4744 | 14 | Bog 14 | 7/10/10 | 45 | AHQN | late |  |
| DSCN4750 | 15 | Tree 15 | 7/11/10 | 32 | AHQN | late |  |
| DSCN4845 | 14 | Bog 14 | 8/11/10 | 1 | AHQN | late |  |
| DSCN4940 | 4 | Lila2 4 | 8/17/10 | 0 | AHQT | early |  |
| DSCN4942 | 1 | Ylva 1 | 8/17/10 | 0 | AHQT | early |  |
| DSCN4944 | 5 | Log 5 | 8/17/10 | 0 | AHQT | early |  |
| DSCN4947 | 6 | Canyon 6 | 8/17/10 | 0 | AHQCanyon | early |  |
| DSCN4949 | 8 | CFH 8 | 8/17/10 | 0 | AHQCanyon | early | overexposed |
| DSCN4955 | 10 | CFL 10 | 8/17/10 | 0 | AHQCanyon | early | overexposed |
| DSCN4956 | 13 | Boulder 13 | 8/17/10 | 0 | AHQCanyon | late | overexposed |
| DSCN4959 | 12 | Bison 12 | 8/17/10 | 0 | AHQCanyon | late | overexposed |
| DSCN5036(1) | 9 | CFM 9 | 9/25/10 | 0 | AHQCanyon | early |  |
| DSCN5039(1) | 10 | CFL 11 | 9/25/10 | 0 | AHQCanyon | early |  |
| DSCN5044(1) | 13 | Bison 12 | 9/25/10 | 0 | AHQCanyon | late |  |
| DSCN5147(1) | 5 | Log 5 | 10/23/10 | 0 | AHQT | early |  |
| DSCN5151 | 6 | Canyon 6 | 10/23/10 | 0 | AHQCanyon | early |  |
| DSCN5159 | 7 | Canyon2 7 | 10/23/10 | 0 | AHQCanyon | early |  |
| DSCN5162 | 8 | CFH 8 | 10/23/10 | 0 | AHQCanyon | early |  |
| DSCN5169 | 9 | CFM 9 | 10/23/10 | 0 | AHQCanyon | early |  |
| DSCN5172 | 10 | CFL 10 | 10/23/10 | 0 | AHQCanyon | early |  |
| DSCN5173 | 13 | Boulder 13 | 10/23/10 | 0 | AHQCanyon | late |  |
| DSCN5176 | 12 | Bison 12 | 10/23/10 | 0 | AHQCanyon | late |  |
| DSCN5316(1) | 12 | Bison 12 | 1/15/11 | 0 | AHQCanyon | late |  |
| DSCN5321(1) | 13 | Boulder 13 | 1/15/11 | 0 | AHQCanyon | late |  |
| DSCN5324(1) | 10 | CFL 10 | 1/15/11 | 0 | AHQCanyon | early |  |
| DSCN5340(1) | 5 | Log 5 | 1/15/11 | 0 | AHQT | early |  |
| DSCN5361 | 1 | Ylva 1 | 1/15/11 | 0 | AHQT | early |  |
| DSCN2222 | 13 | Boulder 13 | 3/12/11 | 0 | AHQCanyon | late |  |
| DSCN2225 | 12 | Bison 12 | 3/12/11 | 0 | AHQCanyon | late |  |
| DSCN2585(1) | 1 | Ylva 1 | 4/15/11 | 37 | AHQT | early |  |
| DSCN2605(1) | 8 | CFH 8 | 4/16/11 | 112 | AHQCanyon | early |  |
| DSCN2608 (1) | 10 | CFL 10 | 4/16/11 | 24 | AHQCanyon | early |  |
| DSCN2610(1) | 13 | Boulder 13 | 4/16/11 | 0 | AHQCanyon | late |  |
| DSCN2613 (1) | 12 | Bison 12 | 4/16/11 | 0 | AHQCanyon | late |  |
| DSCN2737(1) | 12 | Bison 12 | 5/4/11 | 5 | AHQCanyon | late |  |
| DSCN2742 | 13 | Boulder 13 | 5/4/11 | 2 | AHQCanyon | late |  |
| DSCN2743 | 13 | Boulder 13 | 5/4/11 | 2 | AHQCanyon | late |  |
| DSCN2745 | 10 | CFL 10 | 5/4/11 | 108 | AHQCanyon | early |  |
| DSCN2749 | 9 | CFM 9 | 5/4/11 | 155 | AHQCanyon | early |  |
| DSCN2756 | 8 | CFH 8 | 5/4/11 | 471 | AHQCanyon | early |  |
| DSCN2758 | 7 | Canyon2<br>7 | 5/4/11 | 84 | AHQCanyon | early |  |
| DSCN2761 | 6 | Canyon 6 | 5/4/11 | 15 | AHQCanyon | early |  |
| DSCN2764 | 5 | Log 5 | 5/4/11 | 54 | AHQT | early |  |
| DSCN2767 | 4 | Lila2 4 | 5/4/11 | 0 | AHQT | early |  |
| DSCN2775 | 1 | Ylva 1 | 5/4/11 | 73 | AHQT | early |  |

